# Nodamuravirus protein B2 boosts self-amplifying mRNA efficacy by overcoming innate immune barriers

**DOI:** 10.1101/2025.06.27.661928

**Authors:** Raul Y. Sanchez-David, Hoang Duy Le, Josephine Nemegeer, Amanda Gonçalves, Andres Merits, Pierre V. Maillard

## Abstract

Self-amplifying RNA (sa-RNA) technology is a promising strategy for vaccine design, as its intracellular replication boosts transgene expression and provides self-adjuvanticity. However, sa-RNA efficiency is limited by innate immune responses triggered by the presence of intracellular double-stranded RNA (dsRNA). In vertebrates, differentiated cells mainly use type I interferon (IFN) system for protection against viruses, while stem cells rely on IFN-independent mechanisms such as antiviral RNA interference (RNAi). Here, we found that the efficiency of sa-RNAs based on chikungunya virus (CHIKV) or Venezuelan equine encephalitis virus (VEEV) genomes is enhanced when co-expressed in *cis* with the Nodamura virus (NoV) B2 protein, a viral suppressor of RNAi. In stem cells, NoV B2 prevents Dicer-mediated processing of dsRNA, while in somatic cells, it blocks the translation shutdown caused by protein kinase R (PKR), a key effector of the IFN system. Notably, NoV B2 does not interfere with IFN induction and signalling, preserving sa-RNA’s self-adjuvant properties. Mechanistically, NoV B2 sequesters replication-derived dsRNA at the cell periphery, offering a novel strategy to boost sa-RNA efficiency without compromising its immune stimulatory properties.

**Graphical abstract:** 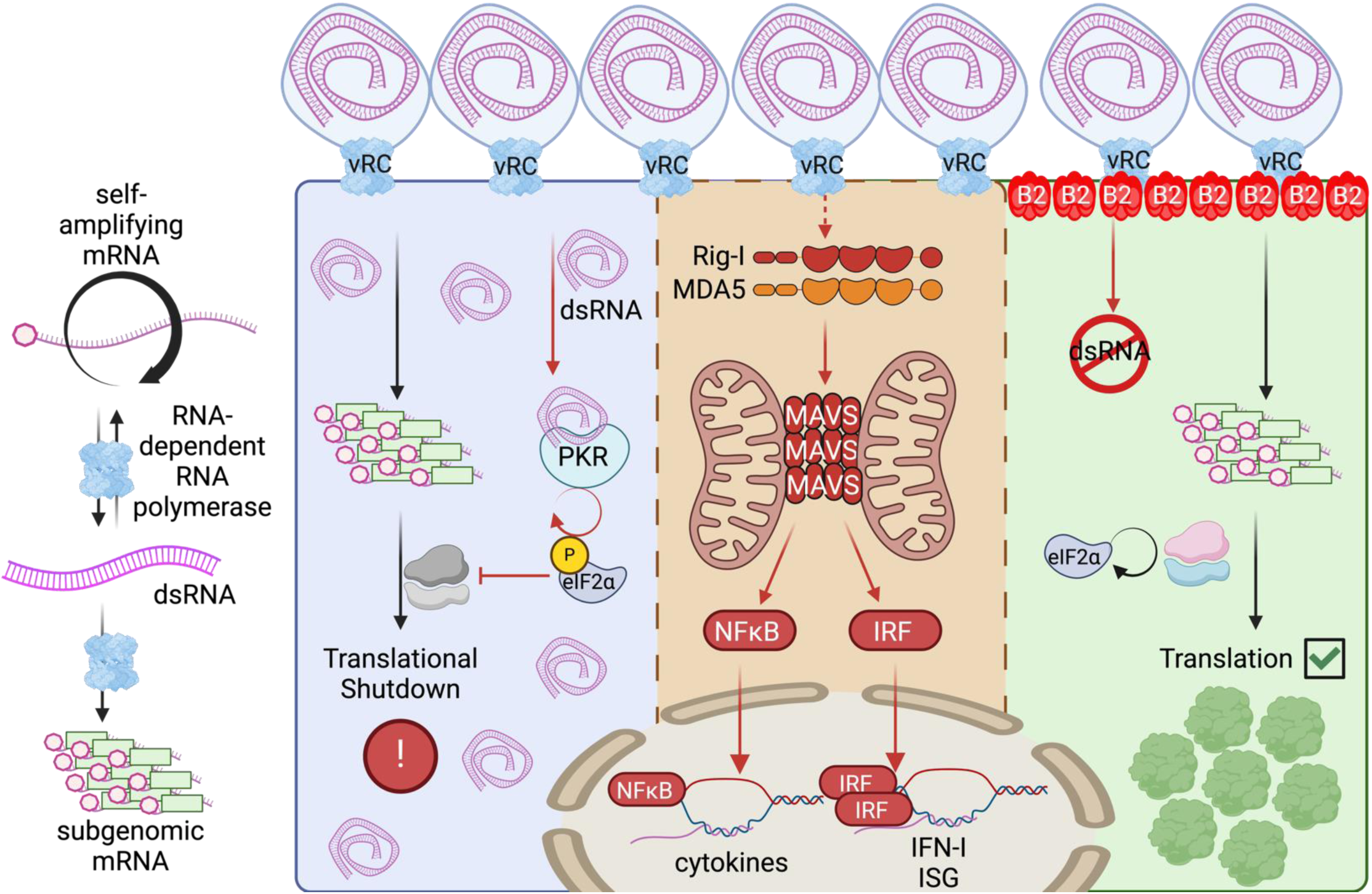

## Introduction

The success of messenger RNA (mRNA)-based COVID-19 vaccines has generated major interest in RNA technologies. Self-amplifying RNA (sa-RNA) holds great promise as a novel strategy for efficient vaccine design. Its ability to replicate within cells enables robust and prolonged antigen expression *in vivo*, thereby allowing lower doses to confer protective immunity compared to non-replicative mRNAs^1–3^.

Sa-RNAs are often derived from alphaviruses, which comprises human pathogens such as chikungunya virus (CHIKV) and Venezuelan equine encephalitis (VEEV)^2,4^. The alphavirus genome is a ∼12 kilobases (kb) long, single-stranded RNA of positive (+) polarity and comprises two open reading frames (ORFs). The 5’ ORF encodes for a polyprotein, which after processing, results in four non-structural proteins (nsP1-4) that form the viral replication complex (vRC). The 3’ ORF is expressed from a subgenomic RNA (sgRNA) produced from an internal promoter located on the negative (-)-sense RNA replication intermediate. The sgRNA encodes for all the structural proteins, which together with newly synthetised genome form new viral particles^4^. In sa-RNAs, all the structural genes are replaced by a gene of interest (GOI) thereby eliminating its capacity to produce viral particles while maintaining its ability to self-amplify^1,2^.

However, sa-RNAs induce a strong innate immune response that enhances their adjuvanticity, yet also hampers their efficiency by activating cell-intrinsic innate antiviral mechanisms detrimental for sa-RNA stability, replication, transcription and/or translation. During self-amplification, vRC generates double-stranded RNA (dsRNA) replication intermediates that trigger innate immune responses such as the interferon (IFN) system^5,6^. In vertebrates, the presence of viral RNAs within the cytosol is sensed by the RIG-I-like receptors (RLRs) that comprise RIG-I (retinoic acid-inducible gene I), MDA5 (melanoma differentiation factor 5) and LGP2 (laboratory of genetics and physiology 2)^5,6^. Upon recognition of agonistic viral RNAs, RIG-I and MDA-5 trigger a signalling cascade via the adaptor protein MAVS (mitochondrial antiviral signalling protein) resulting in the activation of the transcription factors IRF3 and IRF7 (interferon regulatory factor 3 and 7) and NF-κB (nuclear factor kappa-light chain enhance of activated B cells) that eventually induce the expression of type I IFN (IFN-I) and proinflammatory cytokines, respectively^5,6^. Both RIG-I and MDA5-mediated innate immune responses are modulated by LGP2^6–8^. Proinflammatory cytokines initiate inflammation in activated tissues, while IFN-I binds to the IFNAR (IFN-α/β receptor) receptor and signals through the JAK/STAT (Janus Kinase/signal transducer and activator of transcription) pathway to induce hundreds of interferon-stimulated genes (ISGs)^9^. These include cytokines, chemokines and co-stimulatory molecules that promote adaptive immunity^10^. The ability of sa-RNA to generate dsRNA confers strong intrinsic adjuvanticity and enhances immunogenicity compared to non-replicative mRNAs^11–13^. However, many ISGs have antiviral activities that can impair sa-RNA efficiency^9^. For instance, the ISG-encoded serine/threonine kinase PKR (protein kinase R), which plays a central role in the antiviral effects mediated by IFN-I^14^. Albeit being constitutively expressed at steady-state, PKR is upregulated by IFN-I and get activated upon binding to dsRNA of viral or endogenous origin. Upon recognition of dsRNA, PKR phosphorylates the alpha subunit of eukaryotic translation initiation factor 2 alpha (eIF2⍺), thus inhibiting both cellular and viral protein synthesis^14–16^. Supporting the negative impact of the IFN response, sa-RNA protein expression increases when type I IFN signalling is inhibited or in MAVS-deficient mice unable to signal through the RLRs^13^. Similarly, co-expression of some viral IFN antagonists enhances sa-RNA immunogenicity^17^.

The presence of dsRNA in the cytosol elicits another innate antiviral pathway based on RNA interference (RNAi)^18,19^. This pathway acts as the main antiviral defence mechanisms in plants and invertebrates, and recent work shows that it is also active in mammals. In antiviral RNAi, the endoribonuclease enzyme Dicer cleaves viral dsRNAs into viral small interfering RNA (vsiRNAs) of 21 to 23 nucleotides (nt) in length. These vsiRNAs are then loaded into the RNA-induced silencing complex (RISC) composed of Argonaute (Ago) proteins and mediate the sequence-specific cleavage of complementary target viral RNAs^18,19^. Mammalian antiviral RNAi was shown in embryonic and tissue stem cells, which naturally have an attenuated IFN response and express a specific isoform of Dicer called aviD (antiviral Dicer) with increased ability to cleave dsRNA *in vitro* and a heightened antiviral activity^20–23^. In differentiated cells, aviD is expressed at low levels and antiviral RNAi activity is further dampened by IFN-dependent mechanisms^24–27^.

Many viruses encode for viral suppressors of RNAi (VSRs) that inhibit the RNAi pathway and are necessary for efficient virus multiplication and spread^18,28^. Among the best characterised VSRs are the B2 proteins of the *Nodaviridae* family, which includes flock-house virus (FHV) and Nodamura virus (NoV)^18,28^. While FHV infects only insects^29,30^, NoV also causes lethality in suckling mice and pigs^31–33^. B2 proteins homodimerize to bind both synthetic or viral dsRNAs and siRNAs and exert their VSR activity by sequestering them from the RNAi machinery^30,34–37^. Accordingly, NoV mutants lacking B2 or expressing a B2 dsRNA-binding-deficient version (NoV B2^R59Q^) accumulate to lower levels in mouse embryonic stem cells (mESCs) but also in somatic cells (i.e. mouse embryonic fibroblasts) and newborn mice^21,38,39^. These infections are accompanied by the accumulation of canonical vsiRNAs and, in cells, NoV mutant’s replication is rescued upon inactivation of the RNAi pathway.

Given recent evidence that RNAi has antiviral activity in mammalian stem cells and, to a lesser extent, in differentiated cells, we hypothesised that this dsRNA-dependant pathway could limit sa-RNA function. Here we show that *cis*-encoded expression of NoV B2 protein enhances saRNA-driven protein expression from CHIKV and VEEV-based vectors in both mESCs and various somatic cell lines. In mESCs, NoV B2 protects sa-RNA from Dicer-mediated cleavage, while in somatic cells, it prevents PKR-mediated translation block without affecting IFN induction and signalling. Notably, B2 expression causes sa-RNA-derived dsRNA to localise to the cell periphery, suggesting sequestration from the cytosolic PKR pathway.

## Results

### NoV B2 increases sa-RNA-mediated protein expression in mESCs

We engineered a CHIKV-based sa-RNA encoding nsP1-4 and green fluorescent protein (GFP) under the control of a subgenomic promoter. To reduce cytopathic effects and transcriptional shutoff, hallmarks of alphaviruses, we used a mutated version of nsP2 gene (ATL to ERR substitutions at position 674-676) known to prevent these effects while preserving replication^40,41^. To assess the impact of antiviral RNAi on sa-RNA, we replaced the GFP stop codon with a P2A self-cleaving site and the NoV B2 gene (CHIKV-GFP-B2^wt^), a known VSR in mammalian cells^21,38,39^. As a control, we used a version with a stop codon instead of B2 gene (CHIKV-GFP-STOP) (Fig. 1a).

**Figure 1:**
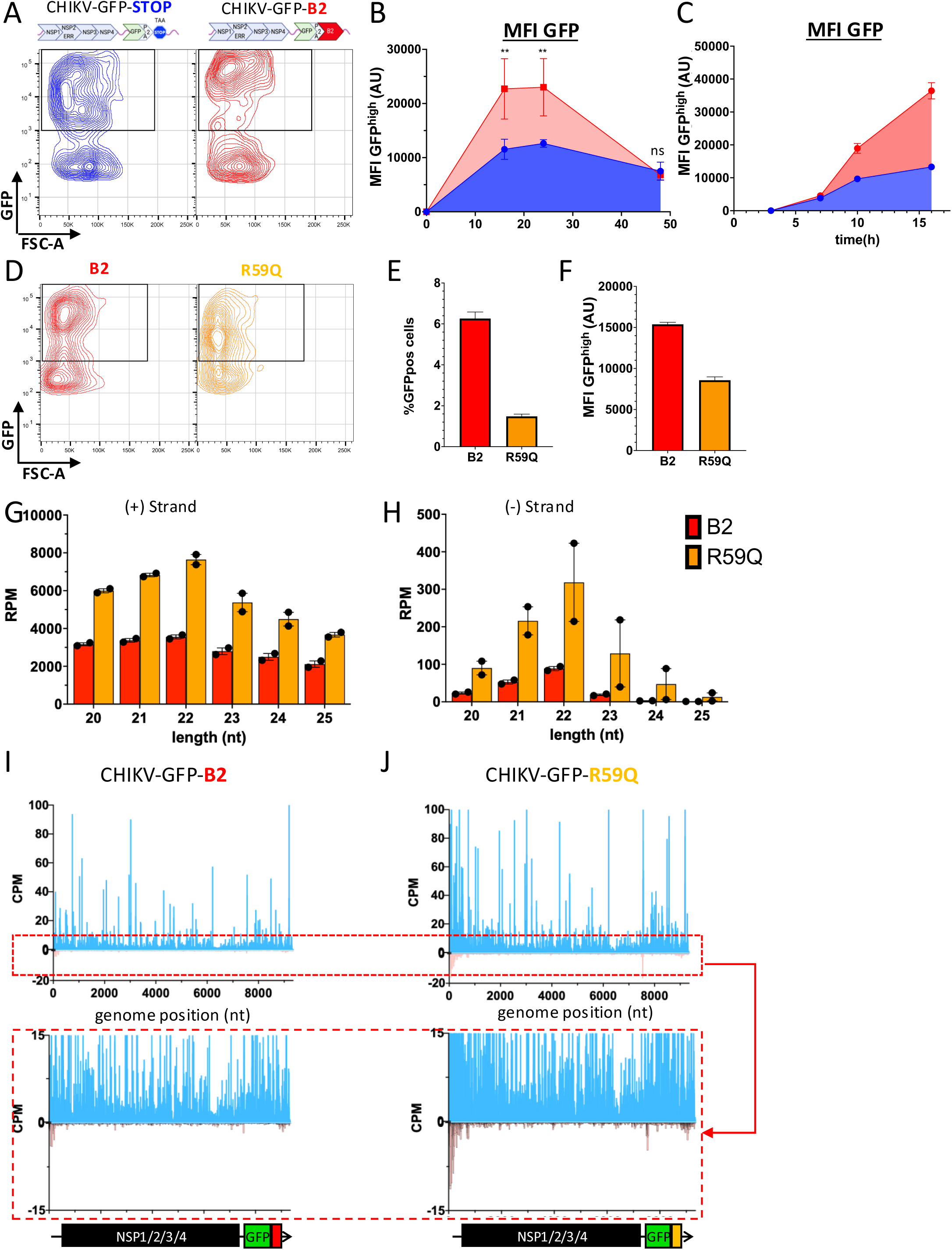
NoV B2 enhances CHIKV-based sa-RNA in mESCs and inhibits the accumulation of ∼22-nt viral small RNAs. **(A)** Representative contour plots of GFP^+^ cells 24 hours after transfection of CHIKV-GFP-STOP (blue) or -B2^wt^ (red) sa-RNA of E14 mESCs. The indicated gate represents GFP^high^ population selected for MFI analysis in (B) and (C). **(B,C)** HeLa cells were transfected with CHIKV-GFP-STOP (blue) or -B2^wt^ (red). Flow cytometry was used to monitor the GFP expression and MFI of GFP^high^ population is represented at (B) 16, 24 and 48 hours after transfection or (C) 3, 7, 10 and 16 hours after transfection. **(D)** Representative contour plots of GFP positive cells from 24 hours after transfection of CHIKV-GFP-B2^wt^ (Red) or B2^R59Q^ mutant (orange) into E14 cells. **(E)** Percentage of GFP^+^ cells and **(F)** MFI GFP^high^ after transfection of indicated sa-RNA. **(G and H)** Distribution of frequencies of small RNAs of different lengths mapping onto the CHIKV-GFP-B2^wt^ (red) or -B2^R59Q^ (orange) vector genome. Each dot corresponds to one biological replicate. **(H and I)** Distribution of 21- and 23-nt reads along the (+) and (–) strands of CHIKV^ERR^ sa-RNA genome after small RNA deep sequencing from GFP^+^-sorted cell population. (I) B2^wt^ or (J) B2^R59Q^. RPM: reads per million normalised to total 15-35 nt long sequences. CPM: counts per million of total 15-35-nt reads.

Mouse embryonic stem cells (mESCs) exhibit a potent RNAi response to synthetic dsRNA and viral infections^21,22,42–44^. To assess the impact of NoV B2 expression, we transfected mESCs with CHIKV-GFP-B2^wt^ or CHIKV-GFP-STOP saRNAs and measured GFP levels over time by flow cytometry. The ability of the vector to amplify the reporter was defined by gating cells with mean fluorescent intensity (MFI) of 10^3^ or higher (GFP^high^, Fig. 1a, Extended Data Fig. 1a). The percentage of GFP-positive (GFP^+^) cells reached a maximum of ∼15% GFP^+^ cells 16 hours post-transfection and were similar between both saRNAs (Extended Data Fig. 1b). Compared to CHIKV-GFP-STOP, the B2^wt^-expressing vector exhibited significantly higher MFI values at 16 and 24 hours post-transfection, while by 48 hours, the difference was no longer detectable (Fig. 1b). Earlier time points showed that GFP expression levels were similar between the two vectors at 3 and 7 hours and only after 10 hours, CHIKV-GFP-B2^wt^ exhibited higher GFP expression (Fig. 1c). To determine whether the B2’s dsRNA-binding activity is necessary for enhanced GFP expression, we introduced an R59Q mutation in NoV B2 (CHIKV-GFP-B2^R59Q^), which impairs dsRNA binding and VSR function^30,36,39,45^. Compared to B2^WT^, transfection with CHIKV-GFP-B2^R59Q^ led to a lower percentage of GFP^+^ cells (Fig. 1e) and reduced MFI within GFP^high^ cells (Fig. 1d-f). These results show that B2, via its dsRNA-binding activity, enhances sa-RNA-mediated gene expression in mESCs.

To assess whether the *cis*-encoded NoV B2 exhibits VSR activity and inhibits Dicer-mediated processing of sa-RNAs, we performed small RNA deep sequencing in GFP^+^ mESCs transfected with CHIKV-GFP-B2^wt^ or CHIKV-GFP-B2^R59Q^. In both cases, 15-35 nucleotides (nt) reads comprised ∼16% of total reads. With B2^WT^ ∼28% mapped to the vector with ∼6% in the 20-25 nt range (Extended Data Fig. 1c). In contrast, B2^R59^ increased 20-25 nt reads to ∼ 13 to 17%. Cells with B2^R59Q^-expressing sa-RNA also showed an accumulation of 22-nt reads from both the viral positive (+) and negative strands (-) (reaching 7’644 reads per million (rpm) and 319 rpm, respectively), characteristic of Dicer activity, whereas B2^WT^ reduced these to 3’555 and 89 rpm, consistent with its VSR function (Fig. 1g-h). We next mapped 21-23-nt viral reads along the viral genome and found that most were derived from the (+)-strand, consistent with the strong (+)-strand bias during alphavirus replication (Fig. 1i,j). However cells with NoV B2^R59Q^-expressing sa-RNA exhibited increased (-)-strand 21-23-nt reads with ∼ 36% originating from the first 500 nucleotides of the vector genome corresponding to regions of dsRNA synthesis known to generate Dicer substrates during alphavirus infections (Fig. 1j)^46^. Thes findings suggest that NoV B2 enhances sa-RNA-mediated gene expression and concomitantly reduces viral siRNA accumulation and that Dicer targeting of sa-RNAs may limit their efficiency in mESCs.

### Enhancement of sa-RNA-mediated protein expression by NoV B2 in somatic cells

Antiviral RNAi activity has also been observed in differentiated mammalian cells infected with wt^46,47^ or VSR-deficient viruses^38,39^. To test NoV B2’s impact on sa-RNA efficiency in cell lines, we compared CHIKV-GFP-STOP and CHIKV-GFP-B2^wt^ in HeLa cells. In CHIKV-GFP-B2^wt^-transfected cells, GFP was robustly enhanced (MFI of 38,000 AU vs 3000 for the control) and this enhancement persisted over time (Fig. 2a-c). We also tested whether NoV B2 could enhance sa-RNA gene expression in *trans*. Co-transfection of Hela cells with CHIKV-GFP-STOP sa-RNA and a plasmid expressing Strep-tagged NoV-B2 (1xSTreP-NoV B2) increased GFP levels compared to cells transfected with an empty plasmid (Extended Data Fig. 2a) confirming that NoV B2 boosts sa-RNA-mediated gene expression both in *cis* or *trans*. Importantly, this enhancement required NoV B2’s dsRNA-binding ability as GFP expression was absent with sa-RNAs encoding NoV B2^R59Q^ (Fig. 2d,f). Cells transfected with CHIKV-GFP-B2^wt^ exhibited higher levels of (+)-sense RNA containing *GFP* or *nsP1* (Fig. 2h), while no increase of (-)-sense RNA was observed (Extended Data Fig. 2b,c). Thus, NoV B2 enhances (+)-strand RNA synthesis and mRNA production through its dsRNA-binding ability.

**Figure 2:**
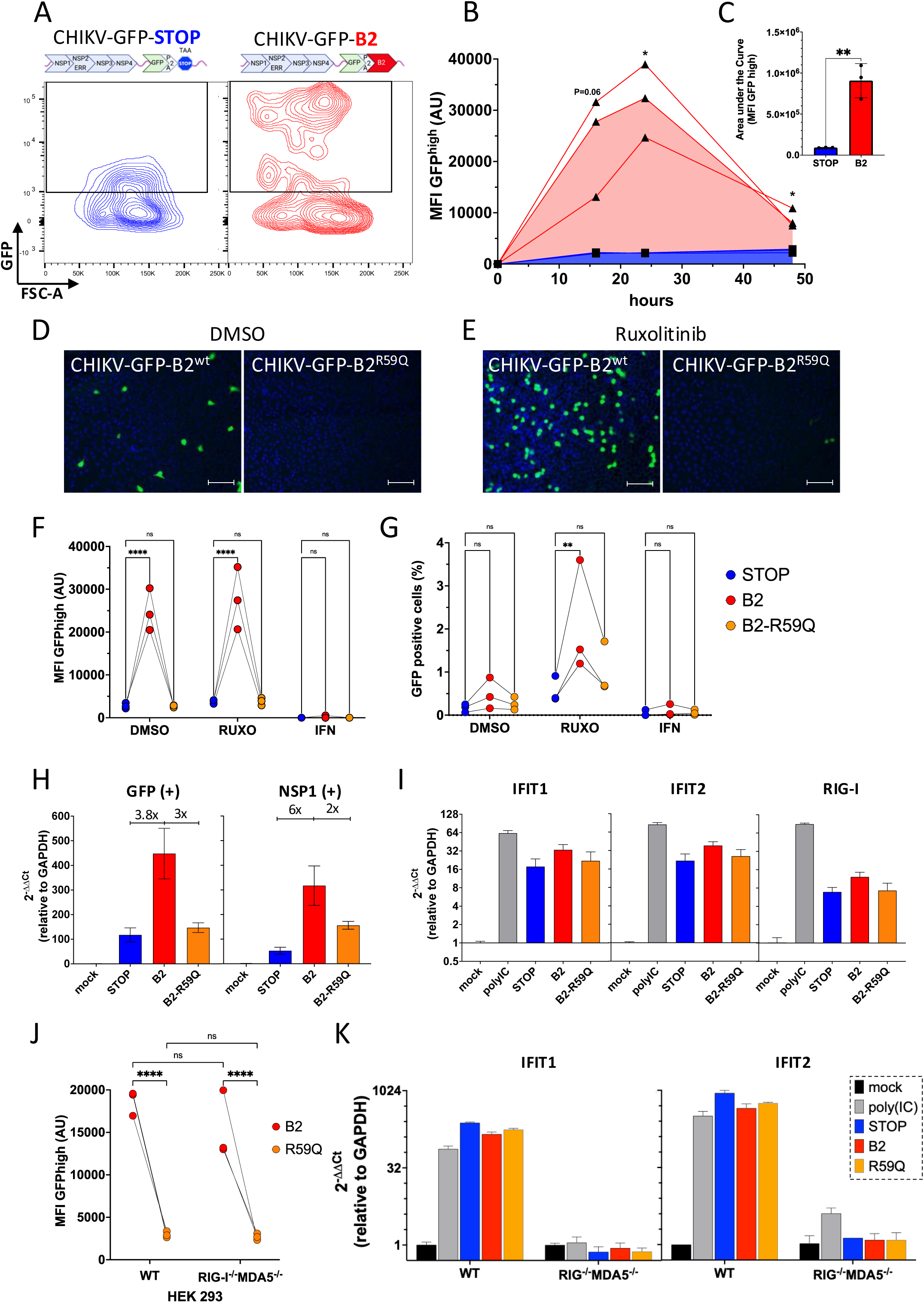
NoV B2 boosts sa-RNA-mediated gene expression in somatic cells independently of type I IFN response. **(A)** Representative contour plots of GFP^+^ cells 24 hours after transfection of CHIKV-GFP-STOP (blue) or -B2^wt^ (red) sa-RNA of HeLa cells. The indicated gate represents GFP^high^ population selected for MFI analysis in (B) and (C). **(B)** HeLa cells were transfected with CHIKV-GFP-STOP (blue) or -B2^wt^ (red). Flow cytometry was used to monitor the GFP expression and MFI of GFP^high^ population at 16, 24 and 48 hours after transfection. **(C)** Area under the curve for GFP^high^ MFI values obtained in (B). **(D and E)** Fluorescence of HeLa cells 24 hours after transfection of CHIKV-GFP-B2^wt^ or -B2^R59Q^ in the presence of DMSO (D) or Ruxotilinib (E). GFP (green) and DAPI (blue). Scale bar: 100 μm. **(F and G)** HeLa cells were transfected with STOP (blue), B2^wt^ (red) or B2^R59Q^ (yellow) expressing sa-RNA in the presence of DMSO, ruxotilinib or recombinant universal IFN-I (IFNα-A/D). 24 hours after transfection, MFI of GFP^high^ (F) or percentage of GFP^+^ cells (G) were plotted. **(H)** RT-qPCR analysis of mRNA specific to GFP (left) or nsP1 mRNA (right) 24 hours after transfection. **(I)** RT-qPCR of ISGs IFIT1 (left), IFIT2 (middle) or RIG-I (right) mRNAs **(J)** GFP^high^ MFI of HEK293 cells WT or RIG-I^-/-^MDA5^-/-^ cells transfected with of CHIKV-GFP-B2^wt^ (red) or -B2^R59Q^ (yellow). **(K)** Cells were treated as in (J) and IFIT1 and IFIT2 mRNAs were analysed by qRT-PCR. Data in all panels are from 3 or 4 independent experiments where each point represents the average of technical replicates. Connected dots correspond to values obtained from the same biological replicate. *P < 0.05, **P < 0.01, ***P < 0.001, ****P < 0.0001 [ratio paired t test (C), two-way ANOVA (other panels)].

Unlike mESCs, somatic cells possess a potent type I IFN system. To determine if NoV B2 enhances sa-RNA-mediated gene by inhibiting IFN signalling, we used ruxolitinib, a JAK1/2 inhibitor blocking IFNAR. GFP levels from CHIKV-GFP-B2^R59Q^ were unchanged by ruxolitinib and NoV B2 still enhanced GFP expression with or without treatment (Fig. 2e,f). However, the overall percentage of GFP+ cells was modulated by the IFN signalling: ruxolitinib treatment increased, while pre-treatment with recombinant IFN-I decreased the percentage of GFP^+^ cells following sa-RNAs transfection regardless of NoV B2 (Fig.2g). These results suggest that while NoV B2 enhances sa-RNA-mediated gene expression independently of signalling, IFN-I influences the overall permissivity of the cell population. Accordingly, no differences in ISGs or cytokine expression (IFIT1, IFIT2, RIG-I and CCL5) were observed following transfection with sa-RNAs encoding either NoV B2^wt^, B2^R59Q^ or no B2, indicating that the IFN signalling pathway is not impacted (Fig.2i and Extented data Fig. 2d). Furthermore, in RIG-I^-/-^ MDA-5^-/-^ cells, which lack RLR-mediated IFN induction (Extended Data Fig. 2e), NoV B2 still enhanced GFP expression (Fig. 2j) without affecting RLRs-dependent ISGs upregulation (Fig. 2k) and resulted in a higher percentage of GFP^+^ cells (Extended Data Fig. 2f), mirroring the effect of ruxolitinib in HeLa cells (Fig. 2g).

To further characterise B2’s effect on sa-RNA, we used THP1-Dual cells (THP1-2R), a human monocytic cell line, which report IFN-I and NF-κB activation via a secreted luciferase and a secreted embryonic alkaline phosphatase (SEAP), respectively (Fig. 3a). CHIKV-GFP-B2^wt^ increased GFP expression compared to CHIKV-GFP-STOP or CHIKV-GFP-B2^R59Q^ (Fig. 3b,c), but didn’t alter IFN-I and NF-κB reporter activity (Fig. 3d, Extended Data Fig.3a). In IFNAR-deficient (*Ifnar2* ^-/-^) THP1-2R, while all three sa-RNAs exhibited a higher percentage of GFP^+^ cells, CHIKV-GFP-B2^wt^ further increased GFP levels compared to controls (Fig. 3c). These results show that while type I IFN limits sa-RNAs efficiency, NoV B2 enhances saRNA-mediated gene expression via an IFN-independent mechanism without affecting ISGs expression.

**Figure 3:**
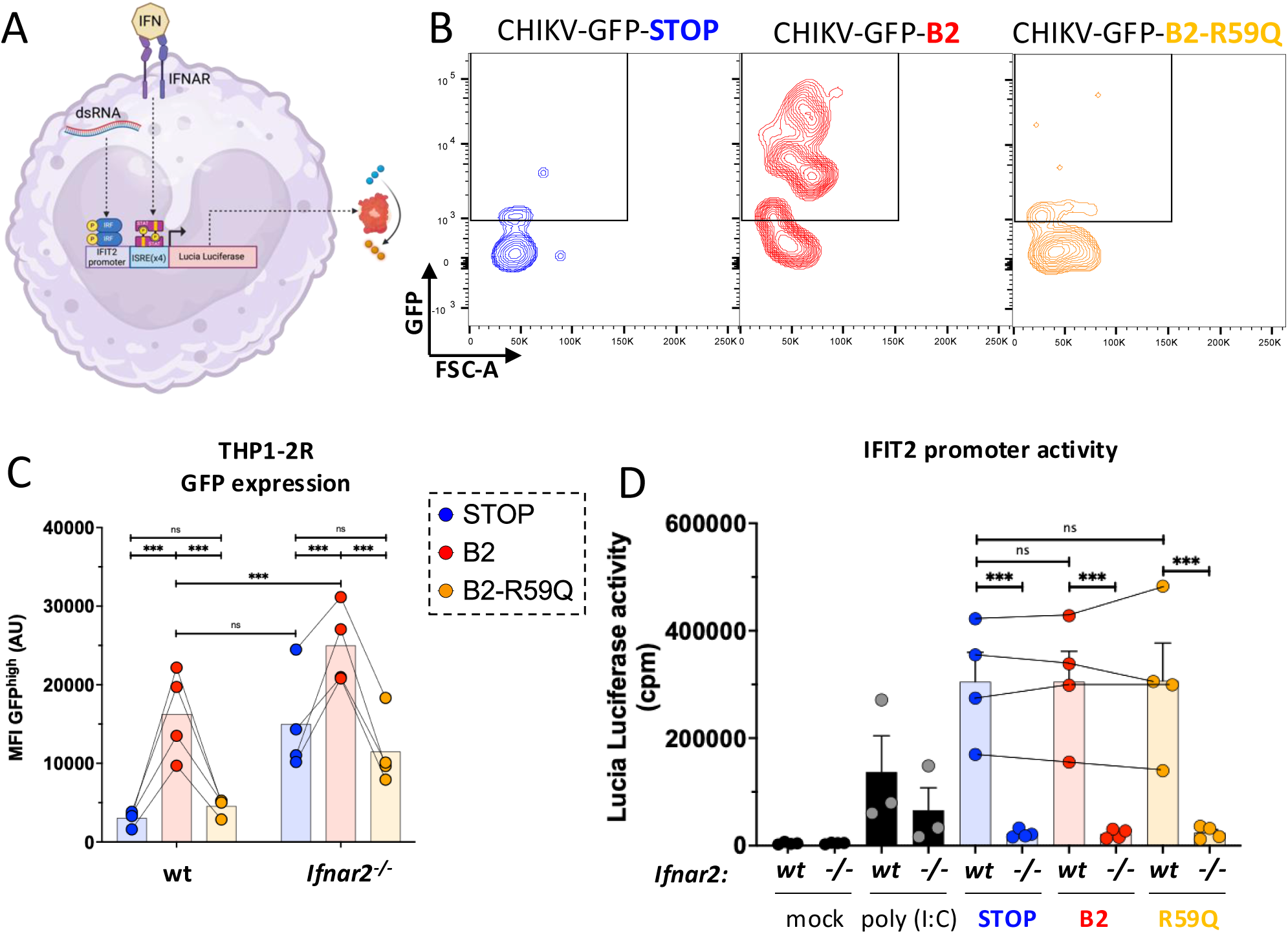
NoV B2 enhances sa-RNA transgene expression in monocytic cells while preserving IFN-I signalling. **(A)** Schematic representation of THP1-2R reporter cell line. Upon activation of the IRF and/or the IFN-I signalling pathway, Lucia Luciferase is secreted. **(B)** Representative contour plots of GFP^+^ THP1-2R reporter cells 24 hours after transfection of CHIKV-GFP-STOP (blue), -B2^wt^ (red) or B2^R59Q^ (orange) sa-RNA. The indicated gate represents GFP^high^ population selected for MFI analysis in (C). **(C)** THP1-2R-wt or IFNAR^-/-^ cells were transfected with CHIKV-GFP-STOP (blue), -B2^wt^ (red) or B2^R59Q^ (orange). Flow cytometry was used to monitor the MFI of GFP^high^ population 24 hours after transfection. **(D)** Immunostimulatory activity of different sa-RNA vectors in THP1-2R-wt or IFNAR^-/-^ cells by measuring Lucia Luciferase in the supernatant. For each experiment in (C), Lucia luciferase activity was measured. Data in all panels are from four independent experiments where each point represents the average of technical replicates. Connected dots correspond to values obtained from the same biological replicate. ***P < 0.001 [two-way ANOVA].

### NoV B2 enhances sa-RNA-mediated gene expression by inhibiting PKR activation in somatic cells

To uncover how NoV B2 enhances sa-RNA-mediated gene expression, we examined whether it protects sa-RNAs from Dicer-mediated cleavage since NoV B2 has known VSR activity not only in stem cells but also in somatic cells^39,45^. We transfected sa-RNAs expressing either NoV B2^wt^, B2^R59Q^ or no B2 into HEK293T cells lacking Dicer (Dicer^-/-^)^48^ or both Dicer and PKR (Dicer^-/-^ PKR ^-/-^) (Fig. 4a, Extended Data Fig. 4a) ^48,49^. The latter was included in our analysis based on previous evidence that Dicer’s antiviral activity against alphaviruses such as SINV and SFV depends on the presence of PKR^50,51^. As in HeLa and THP-1 cells, NoV B2^wt^ expression resulted in a higher MFI in GFP^high^ population in controls HEK293 and HEK293T cells (Fig. 4a). However, GFP expression from CHIKV-GFP-STOP and CHIKV-GFP-B2^R59Q^ remained low in Dicer^-/-^ cells and NoV B2^wt^ -mediated enhancement remained unaffected (Fig. 4a). Strikingly, this restriction was lost in Dicer^-/-^ PKR ^-/-^ cells where GFP levels were similar across all sa-RNAs independently of its expression of a functional NoV B2 (Fig. 4a). We confirmed these results in HeLa PKR^-/-^ cells where the loss of PKR alone was sufficient to relieve restriction thereby eliminating any enhancement from NoV B2 (Fig. 4b). Furthermore. while sa-RNA-mediated expression of NoV B2^R59Q^ was undetectable in HeLa cells, both wild-type and mutant versions of NoV B2 were readily detected in HeLa PKR ^-/-^ cells, demonstrating the loss of restriction (Fig. 4c). Thus, PKR restricts sa-RNAs and the *cis*-encoded NoV B2 alleviate this inhibition.

**Figure 4:**
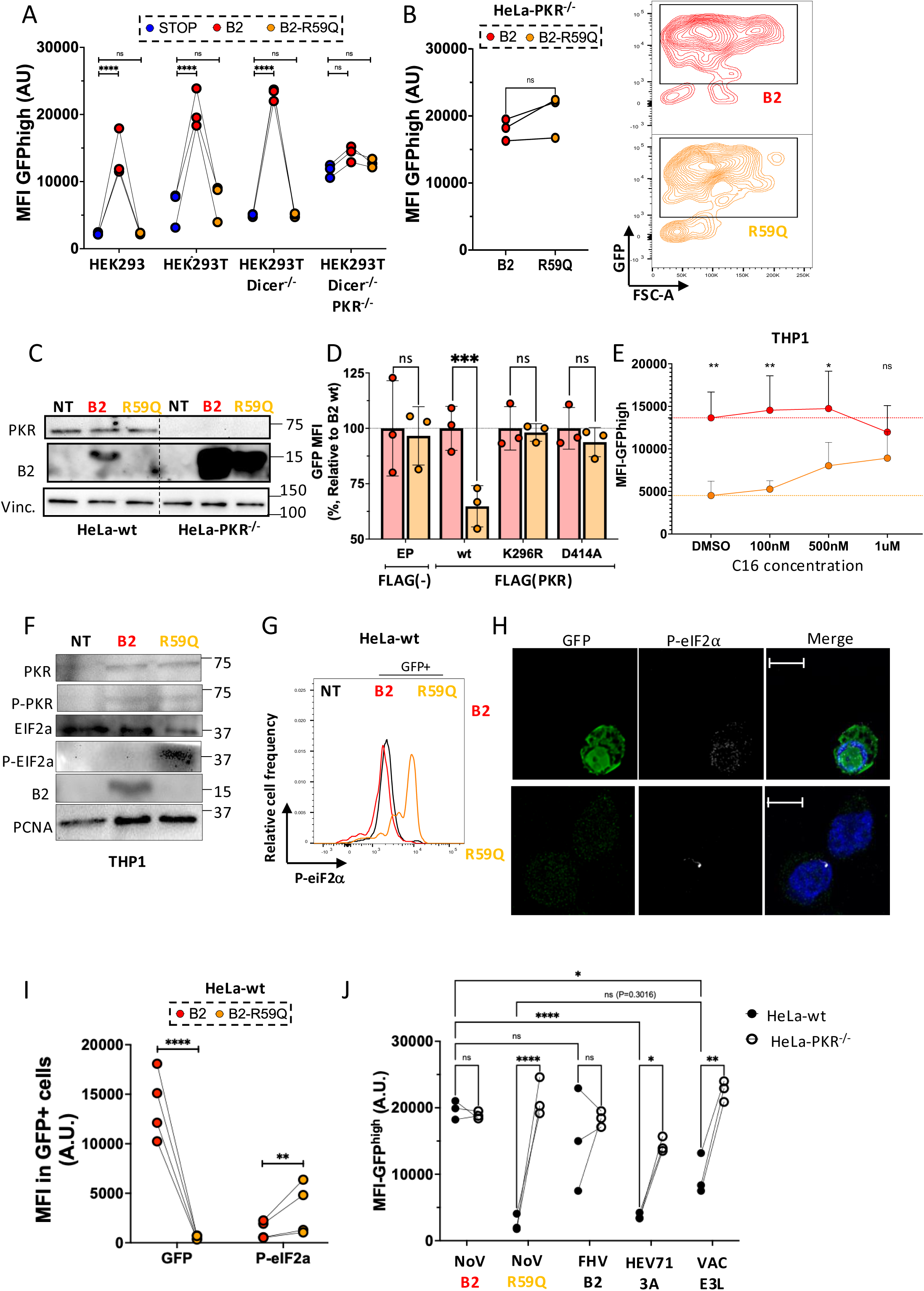
NoV B2 alleviates PKR-mediated restriction of sa-RNA expression by preventing eIF2α phosphorylation. **(A)** HEK293, HEK293T-wt, -Dicer^-/-^ and -Dicer^-/-^ PKR^-/-^ cells were transfected with CHIKV-GFP-STOP (Blue), -B2^wt^ (red) or -B2^R59Q^ (orange). Flow cytometry was used to monitor MFI of GFP^high^ population at 24 hours after transfection. **(B)** HeLa-PKR^-/-^ cells were transfected with CHIKV-GFP-B2^wt^ (red) or -B2^R59Q^ (orange). Representative contour plots of GFP positive cells from 24 hours after transfection were plotted to the right of the histogram. **(C)** Western blot analysis of PKR and B2 expression in HeLa-wt and PKR^-/-^ cells 24 hours after transfection with corresponding CHIKV-GFP-sa-RNA. **(D)** HEK293T-Dicer^-/-^PKR^-/-^ cells were transfected with expression plasmid (EP) or different Flag-tagged PKR encoding plasmids and either CHIKV-GFP-B2^wt^ (red) or -B2^R59Q^ (orange) sa-RNA. 24 hours after transfection, MFI of GFP was measured in the Flag-positive cell population and are plotted as % of GFP MFI relative to B2 expressing vector for each condition. Statistical analyses were calculated based on the MFI-GFP values of three independent experiments. **(E)** THP1-wt cells were transfected with CHIKV-GFP-B2^wt^ (red) or -B2^R59Q^ (orange) in the presence of increasing concentration of C16. GFP-MFI^high^ values of four independent experiments were plotted. **(F)** Western blot analysis of THP1 cells 24 hours after transfection with corresponding CHIKV-GFP-sa-RNA. **(G)** HeLa cells were transfected with CHIKV-GFP-B2^wt^ or -B2^R59Q^ sa-RNA. 24 hours after transfection, GFP positive cells were gated and histogram of P-eIF2⍺ intracellular staining was plotted. **(H)** Confocal microscopy of HeLa cells transfected with corresponding CHIKV-GFP-sa-RNAs for 24 hours and stained for P-eIF2⍺. Scale Bars: 5 μm. **(I)** MFI of GFP-positive HeLa cells for GFP expression and P-eIF2⍺ staining. Data represent the mean values from four independent experiments. Statistical significance was determined using a ratio paired t-test. **(J)** HeLa-wt and PKR^-/-^ cells were transfected for 24 hours with CHIKV-GFP-sa-RNA encoding NoV-B2^wt^, B2^R59Q^ mutant, FHV-B2, HEV71-3A or vaccinia virus E3L. 24 hours after transfection, GFP-MFI^high^ was measured for four independent experiments. *P < 0.05, **P < 0.01, ***P < 0.001, ****P < 0.0001 [two-way ANOVA].

To further characterise PKR restriction, we complemented Dicer^-/-^ PKR ^-/-^ cells with FLAG-tagged constructs encoding either wild-type or mutant versions of PKR. We included PKR^K296R^ (defective in ATP binding, dimerisation and phosphorylation)^52^ and PKR^D414A^, (kinase catalytic mutant)^53^. One day after transfection, cells were transfected with CHIKV-GFP-B2^wt^ or CHIKV-GFP-B2^R59Q^ and GFP levels were assessed in FLAG^+^ cells by flow cytometry (Extended Data Fig. 4b). PKR^wt^ reduced GFP expression from the NoV B2^R59Q^-expressing sa-RNA relative to the B2^wt^-expressing vector, while both PKR mutants failed to restore this restriction indicating that PKR’s kinase activity is essential (Fig. 4d). PKR^wt^ expression was lower than that of PKR^K296R^ and PKR^D414A^ (Extended data Fig. 4c). Yet, phosphorylated PKR (P-PKR) was detected uniquely in PKR^wt^-expressing cells, whereas both PKR mutants were either completely (PKR^K296R^) or partially (PKR^D414^) lacking phosphorylation (Extended Data Fig. 4c).

To confirm that PKR restricts sa-RNAs and that NoV B2 counteract this effect, we transfected THP-1 cells with CHIKV-GFP-B2^wt^ or CHIKV-GFP -B2^R59Q^ in the presence of increasing concentrations of the PKR inhibitor C16, which blocks PKR autophosphorylation^54,55^. In cells transfected with CHIKV-GFP-B2^R59Q^, GFP expression increased in a dose-dependent manner with C16, while no enhancement was observed with CHIKV-GFP-B2^wt^ and GFP levels equalised between the two vectors at 1µM of C16 (Fig. 4e, Extened Data Fig. 4d). Similar results were observed in *Ifnar2* ^-/-^ THP-1 cells, where only 500nM C16 abolished the GFP expression difference between the two sa-RNAs (Extended Data Fig. 4e) likely due to lower PKR expression in the absence of IFN-I signalling. Notably, C16 treatment also increased the percentage of GFP^+^ cells up to eightfold for both sa-RNAs suggesting that blocking steady-state level of PKR activity improves cellular permissivity to sa-RNAs (Supplementary Fig. 4f). These results demonstrate that catalytically active PKR restricts sa-RNA activity and that NoV B2 alleviates this restriction.

We next examined at which step NoV B2 inhibits the PKR pathway. PKR activation involves auto-phosphorylation followed by phosphorylation of eIF2α (P-eIF2α). While Phospho-PKR levels were similar in THP-1 cells transfected with either CHIKV-GFP-B2^wt^ or CHIKV-GFP-B2^R59Q^ (Fig. 4f), no P-eIF2α was detected in CHIKV-GFP-B2-transfected cells where NoV B2 protein was readily expressed (Fig. 4f). Flow cytometry and confocal microscopy confirmed that P-eIF2α levels was detectable in CHIKV-GFP-B2^R59Q^ -transfected cells but absent with CHIKV-GFP-B2^wt^ (Fig. 4g, h). These levels of P-eIF2α inversely correlated with GFP expression from the sa-RNAs (Fig. 4i). Thus, NoV B2 expressed from sa-RNA doesn’t prevent PKR autophosphorylation but blocks eIF2⍺ phosphorylation, bypassing translation inhibition.

We next compared NoV B2 with other VSRs. While Flock house virus B2 (FHV-B2) and human enterovirus 71 (HEV71-3A) have been shown to suppress Dicer activity^30,56,57^, vaccinia virus E3L (VACV-E3L) was shown to inhibit both Dicer and PKR activation^58,59^. Furthermore, VACV-E3L protein expression in *trans* has been shown to enhance vector gene delivery by inhibiting PKR activation and the IFN response^60^. We replaced the NoV B2 in CHIKV-GFP-B2 with FHV-B2, HEV71-3A or VACV-E3L and tested sa-RNA efficiency in both wild-type HeLa and PKR ^-/-^ cells (Fig. 4j). FHV-B2 enhanced sa-RNA activity similarly to NoV-B2, while HEV71-3A did not counteract this restriction. However, VACV-E3L modestly improved GFP levels compared to NoV B2^R59Q^ and VACV-E3L-expressing sa-RNAs exhibited lower activity in wild-type relative to PKR ^-/-^ cells, suggesting only partial inhibition of PKR. Thus, among the tested VSRs, the *cis*-expression of NoV-B2 and FHV-B2 most effectively alleviate PKR-mediated restriction of sa-RNAs.

### *Cis*-expression of NoV B2 results in a change of dsRNA spatial distribution within the cells

To further investigate the mechanism by which B2 enhances sa-RNA-mediated gene expression, we analysed its subcellular localization by confocal microscopy. A functional N-terminal STrEP-tagged NoV B2 protein was expressed from a sa-RNA-GFP and was detected predominantly at the cell periphery (Fig. 5a). Since alphavirus RNA replication complex (vRC) form at the plasma membrane, we compared B2 localisation to that of nsP1, a membrane-ancored vRC component that facilitates the spherules formation, invaginations that harbour viral dsRNAintermediates ^61–65^. While nsP1 was localised as expected at the cell periphery, B2 was found adjacent to, but not overlapping with, nsP1 (Fig. 5a). We next analysed nsP2, a vRC component located on the cytoplasmic side of the spherules^63^. nsP2 exhibited diffused cytoplasmic distribution with partial colocalisation with B2 at the cell periphery (Fig. 5b). We then analysed the localisation of STrEP-tagged B2^R59Q^. As CHIKV sa-RNAs encoding a non-functional NoV B2 are restricted in HeLa cells, we assessed the mutant distribution in HeLa PKR ^-/-^ cells. While NoV B2^wt^ still localised to the periphery, NoV B2^R59Q^ was diffusely distributed throughout the cytoplasm (Fig. 5c, d and Extended Data Fig. 5c). Similar results were obtained in *Ifnar2* ^-/-^ THP-1 cells where NoV B2^wt^ was at the cell periphery showing no overlap with GFP, while NoV B2^R59Q^ colocalised more with cytoplasmic GFP (Extended Data Fig. 5a, b). Given the importance of NoV B2’s dsRNA-binding activity for its localisation, we assessed dsRNA distribution in HeLa PKR ^-/-^ cells transfected with CHIKV sa-RNA encoding either STrEP-B2^wt^ or -B2^R59Q^. Without functional B2, dsRNA was dispersed throughout the cytoplasm (Fig. 5c, Extended Data Fig. 5c). In contrast, B2^wt^-expressing cells had reduced cytoplasmic dsRNA and enhanced accumulation at the cell periphery (Fig. 5d, Extended Data Fig. 5c). Orthogonal views showed NoV B2 localised just beneath dsRNA foci, without direct overlap (Fig. 5d).

**Figure 5:**
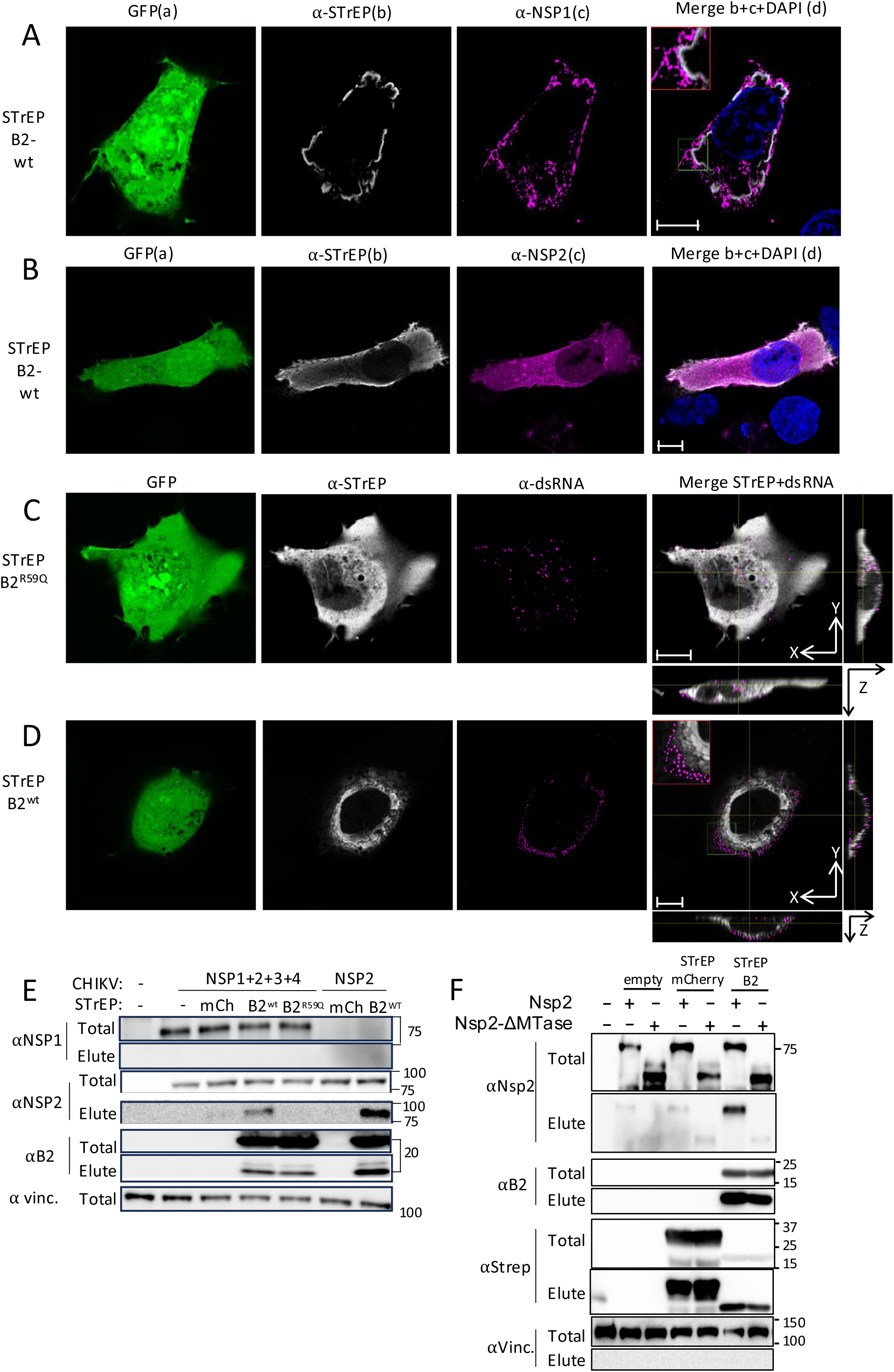
NoV B2 redistributes dsRNA to the cell periphery and interacts with nsP2 in a dsRNA-binding-dependent manner. **(A, B)** Confocal microscopy image of HeLa cells transfected with CHIKV-GFP-STrEP-B2 sa-RNA stained for nucleus (blue), STrEP-tag (white) and nsP1(A) or nsP2 (B) (magenta). Red square in panel (A-d) corresponds to a magnification of the green square in the same image. **(C,D)** Confocal microscopy images of HeLa-PKR^-/-^ cells transfected with CHIKV sa-RNA encoding GFP-STrEP-B2^wt^ (C) or GFP-STrEP-B2^R59Q^ (D). Cells were stained for STrEP-tag (white) and dsRNA (magenta). Red square in panels (F-d, C-d) corresponds to a magnification of the green square in the same image. Panels (F-d) and (C-d) show orthogonal views of the distribution of STrEP and dsRNA staining. Scale Bars: 10 μm. **(E)** The indicated STrEP-tagged constructs (STrEP-mCherry, -NoV-B2^wt^ or -NoV-B2^R59Q^) were co-transfected with four constructs encoding nsP1, nsP2, nsP3 and nsP4 or nsP2 alone in HEK293T cells followed by STrEP immunoprecipitation, SDS-PAGE and western blot using the indicated antibodies. **(F)** Same as in (E), but the indicated STrEP-tagged constructs were co-transfected with constructs expressing either full-length (nsP2) or MTase-like domain truncation mutant of nsP2 (nsP2-ΔMTase) followed by STrEP immunoprecipation.

Given B2’s localisation beneath the vRC, we tested whether it interacts with CHIKV replication components. Co-transfection of STrEP-tagged B2^wt^ or B2^R59Q^ with CHIKV nsP1-4 or nsP2 alone revealed that NoV B2^wt^ specifically co-precipitated with nsP2, and not nsP1, and that this interaction was independent of other vRC components (Fig. 5e). In contrast, NoV B2^R59Q^ was no longer able to interact with nsP2 (Fig. 5e). Deletion of the C-terminal methyltransferase (MTase)-like domain of nsP2 also abolished binding to NoV B2 (Fig. 5f). These results indicate that NoV-B2’s localisation near the vRC at the cell periphery and its ability to bind nsP2 and/or dsRNA are critical for this subcellular distribution and ability to alleviate PKR restriction.

### NoV B2 enhances gene expression from VEEV-based sa-RNAs

To assess whether NoV B2 also boosts gene expression from vectors derived from other alphavirus genomes, we examined its impact on Venezuelan equine encephalitis virus (VEEV)-based sa-RNAs that are also currently used for vaccine studies^2^. VEEV-based vectors encoding GFP and either NoV B2^wt^ or B2^R59Q^ were tested in various cell types. VEEV-based vectors were more efficient and displayed higher percentage of GFP^+^ cells in HEK293, HEK293T and THP1 cells compared to CHIKV-based vectors (Extended Dat Fig. 6a). HeLa cells remained the most restrictive cells for both vectors. In HeLa cells, the *cis*-expression of NoV B2 from VEEV sa-RNA increased the MFI of the GFP^high^ population, an effect that was lost in PKR ^-/-^ cells, confirming PKR as the main restriction factor (Fig 6a,b). Confocal microscopy showed that NoV B2^wt^ localised to the cell periphery with dsRNA foci forming an outer layer (Fig. 6c, Extended Data Fig. 6b), whereas in NoV B2^R59Q^-expressing cells both B2^R59Q^ dsRNA remained diffuse in the cytoplasm, similar to observations with CHIKV vectors (Fig. 6d, Extended Data Fig. 6b). Thus, VEEV-derived vectors, like CHIKV-based sa-RNAs, are restricted by PKR, and NoV B2 can alleviate this restriction by spatially redistributing the dsRNA.

**Figure 6:**
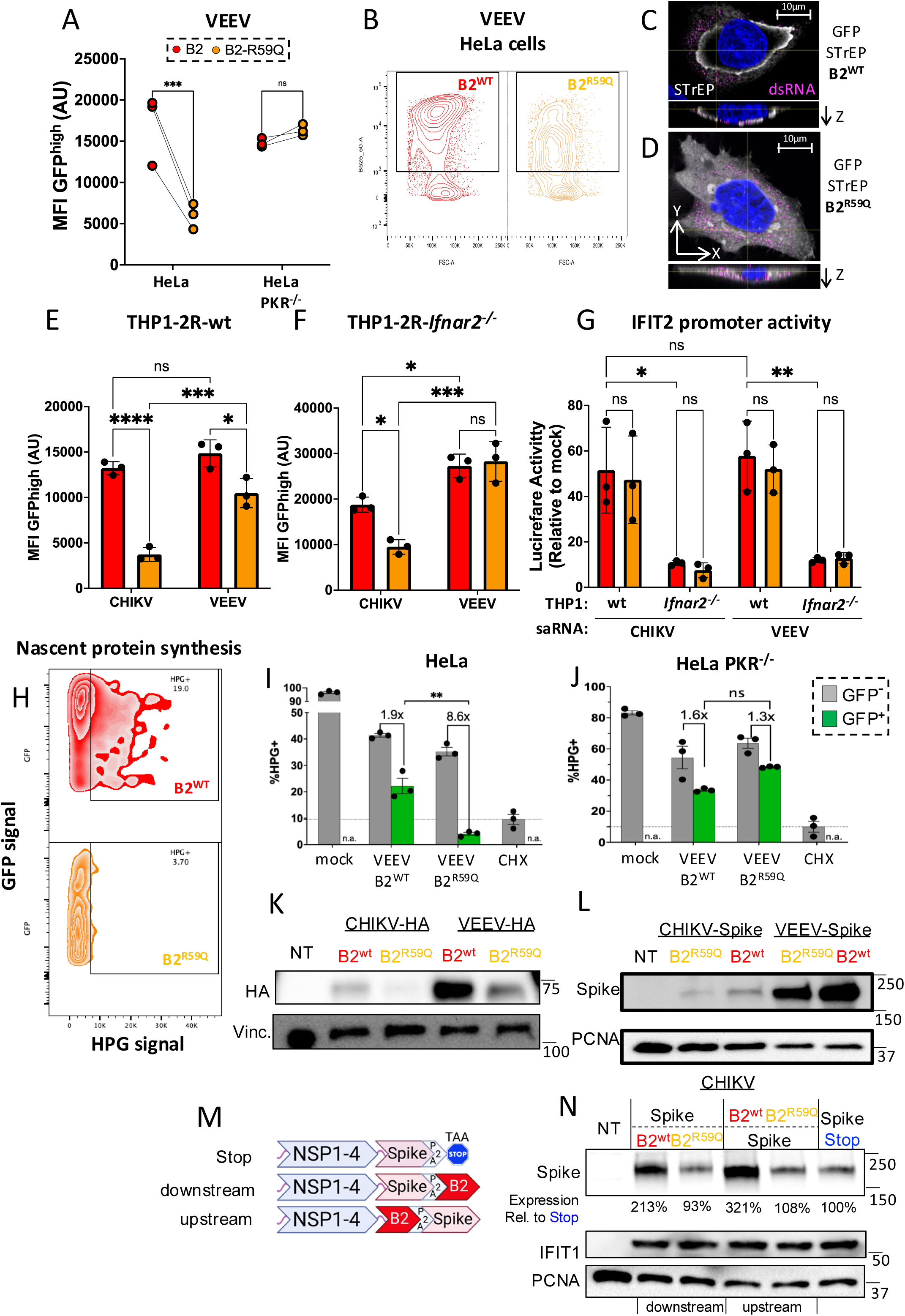
NoV B2 boosts expression from VEEV-based sa-RNA by counteracting PKR-dependent translational inhibition without altering IFN responses. **(A)** HeLa-WT and HeLa-PKR^-/-^ cells were transfected with VEEV-GFP-B2^wt^ (red) or -B2^R59Q^ (orange). Flow cytometry was used to monitor MFI of GFP^high^ population 24 hours after transfection. **(B)** Representative contour plots of GFP^+^ cells 24 hours after transfection with VEEV-GFP-B2^wt^ and -B2^R59Q^ vectors. **(C,D)** Confocal microscopy images of HeLa-wt cells transfected with VEEV-GFP-STrEP-B2^wt^ (C) or -STrEP-B2^R59Q^ (D). Scale Bars: 10 μm. Cells were stained for STrEP-tag (white) and dsRNA (magenta) and DAPI (blue). Both images present orthogonal views of the distribution of STrEP and dsRNA staining. **(E)** THP1-2R-wt or **(F)** IFNAR^-/-^ cells were transfected with CHIKV^ERR^-GFP- or VEEV-GFP-B2^wt^ (red) or - B2^R59Q^ (orange). **(G)** Immunostimulatory activity of different sa-RNA vectors in THP1-2R-wt or -IFNAR^-/-^ cells by measuring Lucia Luciferase in the supernatants of (E) and (F). **(H-J)** Flow cytometry was used to monitor nascent protein synthesis in HeLa-wt (H and I) or PKR^-/-^ (J) cells transfected with VEEV-derived sa-RNA expressing GFP-B2^wt^ or GFP-B2^R59Q^. (H) Representative contour plot of GFP^+^ cells after HPG labelling. **(I and J)** Comparative analysis of HPG incorporation in GFP^+^ and GFP^-^ cells in different conditions. **(K)** Western blot analysis of IAV-HA protein expression in HEK293T.17 cells 24 hours after transfection of CHIKV-HA- or VEEV-HA-B2^wt^ or -B2^R59Q^ expressing sa-RNA. **(L)** Same as (K) but for SARS-CoV-2 Spike prefusion protein expressed by the indicated sa-RNA. (M) Same as (L), but in HeLa-wt cells with vectors expressing NoV-B2^wt^ or NoV-B2^R59Q^ upstream or downstream of SARS-CoV-2 Spike prefusion protein. Data in all FACS panels are from three independent experiments where each point represents the average of technical replicates. *P < 0.05, **P < 0.01, ***P < 0.001, ****P < 0.0001 [two-way ANOVA].

In THP1 cells, VEEV-B2^R59Q^ vectors displayed significantly higher activity than CHIKV-B2^R59Q^, suggesting that VEEV is naturally less sensitive to PKR activity in THP1 cells (Fig. 6e). Nonetheless, NoV B2 further enhanced gene expression from VEEV-based sa-RNAs and its expression abolished the differences in GFP expression observed between CHIKV and VEEV-based vectors (Fig. 6e). This suggests that PKR inhibition by NoV B2 overcomes inherent differences in vector sensitivity. Importantly, as previously observed with CHIKV-based sa-RNA, there was a similar activation of the IFN-I response in cells transfected with VEEV expressing NoV B2^wt^ or B2^R59Q^, but the B2-mediated enhancement of GFP was lost in *Ifnar2* ^-/-^ THP1-2R (Fig.6f,g), suggesting that the boosted expression of PKR upon IFN-I signalling amplification is responsible for the restriction in the VEEV system.

To explore the broader impact of NoV B2 on the global host translation, we measured nascent protein synthesis using the incorporation of L-homopropargylglycine (HPG), a methionine analogue which can stain cells by click chemistry^66^. VEEV sa-RNA transfection induced a strong reduction in protein synthesis across the entire cell population (Extended Data Fig. 6c). HPG incorporation was reduced by more than 50% in the bulk population, reaching levels comparable to cells treated with cycloheximide (CHX), a widely used translation inhibitor (Extended Data Fig. 6c). To differentiate between sa-RNA-active and inactive cells, we measured HPG incorporation specifically in the GFP⁺ population (Extended Data Fig. 6d). In GFP^+^ cells, NoV B2^R59Q^ caused near-complete translational shutdown, showing an 8.6-fold reduction compared to GFP⁻ cells, while NoV B2^WT^ partially restored protein synthesis (Fig. 6h, i). This rescue was PKR-dependent as it was absent in HeLa-PKR⁻/⁻ cells, (Fig. 6j). Thus, the inclusion of B2^wt^ not only enhances transgene expression but also inhibit the global translational shutdown of host mRNAs, promoting a more permissive environment for protein synthesis. Finally, NoV B2^WT^ enhanced expression of clinically relevant antigens such as Influenza A virus PR8 hemagglutinin (HA) or the prefusion form of the SARS-CoV-2 Spike protein from both CHIKV and VEEV vectors (Fig. 6k, l). In addition, we examined how the position of the B2 gene affects antigen expression in CHIKV-based vectors in HeLa cells. Placing NoV B2^WT^ downstream of the Spike gene resulted in a 2-fold increase in protein expression, while positioning it upstream yielded a 3-fold increase (Fig. 6m, n). In contrast, B2^R59Q^ had no effect on antigen expression regardless of position. Consistent with our previous observations, NoV B2^WT^ did not alter the type I IFN response since the expression of IFIT1, a well-established ISG, remained unchanged across all five tested vectors (Fig.6n).

## Discussion

sa-RNA boosts antigen expression and acts as a self-adjuvant, but cytosolic dsRNA generated during replication activate innate antiviral responses that restrict its efficiency. Here we show that *cis*-expression of NoV B2 protein enhances sa-RNA-mediated protein expression from CHIKV or VEEV-based vectors, two of the most promising backbones for vaccination^67^, in both stem cells and somatic cells.

In mESCs, CHIKV sa-RNAs triggered accumulation of 21-23-nt small RNAs primarily mapping to the 5’end of the vector’s genome and the subgenomic transcription start site. These reads derived from both strands, indicative of Dicer processing of replication-derived dsRNA replication intermediates. These findings are consistent with antiviral RNAi, previously shown to be active in stem cells, that lacks an IFN response and instead rely on alternative defence mechanism for viral defence^18,68–71^. Similar patterns of siRNAs accumulation have been shown in infections with viruses restricted by antiviral RNAi, such as Semliki Forest virus, Sindbis virus, Zika virus and NoV^30,39,46,72,73^. The abundance of siRNA detected in CHIKV-based sa-RNAs might be modulated by VSRs as CHIKV nsP2 and nsP3 have been shown to inhibit shRNA and siRNA-mediated RNAi, though their VSR activity during infection remains to be addressed^74^. Consistent with this, *cis*-expression of NoV B2, a well-established VSR ^21,30,38,39^, reduced 21-23-nt viral reads accumulation and enhanced sa-RNA-mediated protein expression, suggesting that bypassing antiviral RNAi can improve sa-RNA efficiency. These findings in mESCs may exend to other stem cells, including human induced pluripotent stem cells (hiPSCs) and adult stem cells, that might have a potent RNAi response due to their expression of aviD, a Dicer isoform with enhanced dsRNA processing activity^22^. Further work should address whether other pathways might contribute to the overall restriction of sa-RNA in stem cells.

Antiviral RNAi activity declines upon cellular differentiation with reduced viral siRNA accumulation in infected cells^21^. This has been linked to an antagonism between antiviral RNAi and the IFN system and to low levels of aviD expression in differentiated cells ^22,24–26^. Yet, some studies showed an antiviral RNAi activity during infections of somatic cells with virulent or VSR-deficient viruses suggesting that, despite the action of inhibitory mechanisms, the RNAi response is still active in differentiated cells^39,46,47,56,75^.

We found that NoV B2 enhanced sa-RNA-driven gene expression in three IFN-competent somatic cells (i.e. HeLa, THP-1 and HEK293 cells) without suppressing ISGs induction. While the exact ligand of sa-RNA recognised by RLRs needs to be identified, parental CHIKV infections would suggest that the 3’UTR of the genomic and subgenomic RNA would still harbour immunostimulatory activity^76^. Our findings align with previous studies showing that NoV B2 suppresses RNAi but not the IFN-I response. Indeed, B2 expression from a NoV replicon had minimal impact on ISGs or IFN-β and infections of mouse embryonic fibroblasts with NoV (i.e. naturally encoding B2) still triggered a robust RIG-I-mediated IFN response^39,77^. Interestingly, previous studies found that *cis*-expression of the V protein from PIV-5 (parainfluenza virus-5) or ORF4a from MERS-CoV (Middle East respiratory syndrome coronavirus) enhanced sa-RNA-mediated protein expression by suppressing the IFN-I pathway^17^. In contrast, our findings indicate that NoV B2 uniquely boosts sa-RNA-gene expression without impairing IFN signalling, potentially preserving the vector’s self-adjuvant properties.

Our analysis further shows that NoV B2 enhances sa-RNA-mediated protein expression in somatic cells through a mechanism independent of Dicer. Instead, CHIKV and VEEV sa-RNAs are restricted by PKR in various cell lines and NoV B2 confer resistance to this inhibition. Complementation experiments confirmed that this restriction was dependent on PKR activity and that *cis*-expression of NoV B2 prevented the phosphorylation of eIF2α, a key PKR substrate, thereby preventing translational shutdown. This function requires NoV B2’s dsRNA-binding activity as NoV B2^R59Q^ failed to inhibit eIF2α phosphorylation. While NoV B2’s ability to bind dsRNA and shield it from Dicer’s activity is well-documented, our findings reveal an additional role in inhibiting the PKR pathway when expressed from CHIKV or VEEV sa-RNAs. Whether this PKR-suppressive function of B2 also occurs during NoV infections in mammalian cells remains to be determined. Interestingly, previous studies showed an interplay between PKR and Dicer. First, PKR was shown to interact with Dicer’s helicase domain to prevent aberrant NF-κB pathway activation^50,51^. Secondly, Dicer was proposed to regulate PKR activation in mESCs, yet the mechanism remains unclear^78^. Finally, both Dicer and PKR share cofactors, including TRBP and PACT, which modulate their activity^79^. Further studies are needed to determine whether the interactions between Dicer, PKR, TRBP and/or PACT influence NoV B2’s suppressive functions.

Consistent with a previous study^60^, VACV-E3L also enhanced sa-RNA activity in our system, yet not as strongly as NoV B2, suggesting mechanistic differences. Like NoV B2, VACV-E3L is a dsRNA-binding protein and is known to sequester dsRNA from its recognition by PKR^60,80^.Yet, NoV B2’s ability to boost sa-RNA may additionally encompass a specific compatibility with alphaviruses’ replication complex. Alphavirus forms spherules at the plasma membranes, where nsP1, assembling into a 12-mer ring, anchors the complex at the plasma membrane and nsP4 synthesizes the (–)-strand viral antigenome which serves as a template for genomic and subgenomic mRNAs production and is the primary source of viral dsRNA^63,81,82^. NoV encodes multifunctional viral replicase Protein A that exhibits striking parallels with the alphavirus vRC with the two N-terminal domains of Protein A structurally similar to alphavirus nsP1^62,83^. Like nsP1, NoV Protein A multimerises at the outer mitochondrial membrane (OMM) and forms similar 12-mer ring-like structures ^84,85^. FHV B2 was shown to interact with Protein A in addition to binding viral dsRNA in FHV-infected insect cells^30^. It was therefore proposed that B2 functions as a structural component of the FHV vRC at the OMM, ideally positioned to sequester dsRNA from Dicer.

Our data suggest that NoV B2, when expressed from CHIKV and VEEV sa-RNA in somatic cells, functions similarly, localising beneath nsP1 at the cell periphery and interacting with nsP2. Notably, this interaction occurs via a conserved MTase-like domain with nsP2, a structurally defined but functionally poorly characterised domain found across alphaviruses^86^. Although this domain lacks demonstrable methyltransferase activity, it shares structural homology with the N-terminal MTAse-GTPase region of nsP1 as well as with the MTAse-like domain found in nodavirus Protein A^83,86^. The ability of NoV B2 to bind nsP2 may facilitate its integration into the vRC ideally positioning it adjacent to the alphavirus replication machinery to shield dsRNA from the PKR pathway. Supporting this, we observed enrichment of dsRNA at the cell periphery in cells expressing NoV B2^wt^. This spatial positioning may stabilise dsRNA-containing spherules at the plasma membrane and limit access to PKR and its main substrates, eiF2α. Whether similar mechanisms operate during natural NoV or FHV infections remained to be addressed. though detection of spatial redistribution of NoV/FHV dsRNA at the mitochondrial membrane might be challenging given that this organelle is scattered throughout the cell’s cytoplasm.

Given the success of sa-RNA vaccines at low doses in humans, our findings that NoV B2 enhances sa-RNA efficiency in somatic cells without disrupting innate immune signalling offers a strategy to improve immunogenicity while preserving self-adjuvanticity. Furthermore, since sa-RNA delivery of reprogramming factors was shown as an efficient approach to generate human iPSCs, NoV B2 may also enhance sa-RNA-based regenerative therapies^87^. In addition, our results reveal that sa-RNAs trigger a global translation shutdown underscoring the cellular stress they impose. Co-expression of NoV B2 reduces this global response and thereby enhances the cell’s capacity for protein synthesis. Since sa-RNAs have been linked to liver toxicity^88^, future studies should evaluate the impact of NoV B2 on sa-RNA-associated cytotoxicity both *in cellulo* and *in vivo*.

## Acknowledgements

We thank all members of the Maillard lab for helpful discussions, the Blizard Institute Genome Centre, the Flow Cytometry Facility, the Blizard Advanced Light Microscopy Facility for technical assistance. We further thanks Dr Monika Hamilton, Dr Ben Milborne and all the Queen Mary Innovation team for their advice; Prof. Adam Gaballe, Prof. Bryan R. Cullen, Dr Annemarthe Van der Veen for their kind gift of cells; Prof. Shou-Wei Ding for his kind gift of antibodies. This work was supported by a UK Research and Innovation Future Leaders Fellowships (FLF) (MR/S034498) and a Barts Charity Rising Stars Programme grant (MGU0459) awarded to P.V.M.

## Author contributions

R.Y.S.D and P.V.M. designed the experiments, analysed data and wrote the manuscript. R.Y.S.D conducted experiments with assistance from J.N and H.L. A.M. provided key reagents and expertise. P.V.M supervised the project. All authors reviewed the manuscript.

---

**Extended Data Figure 1.**
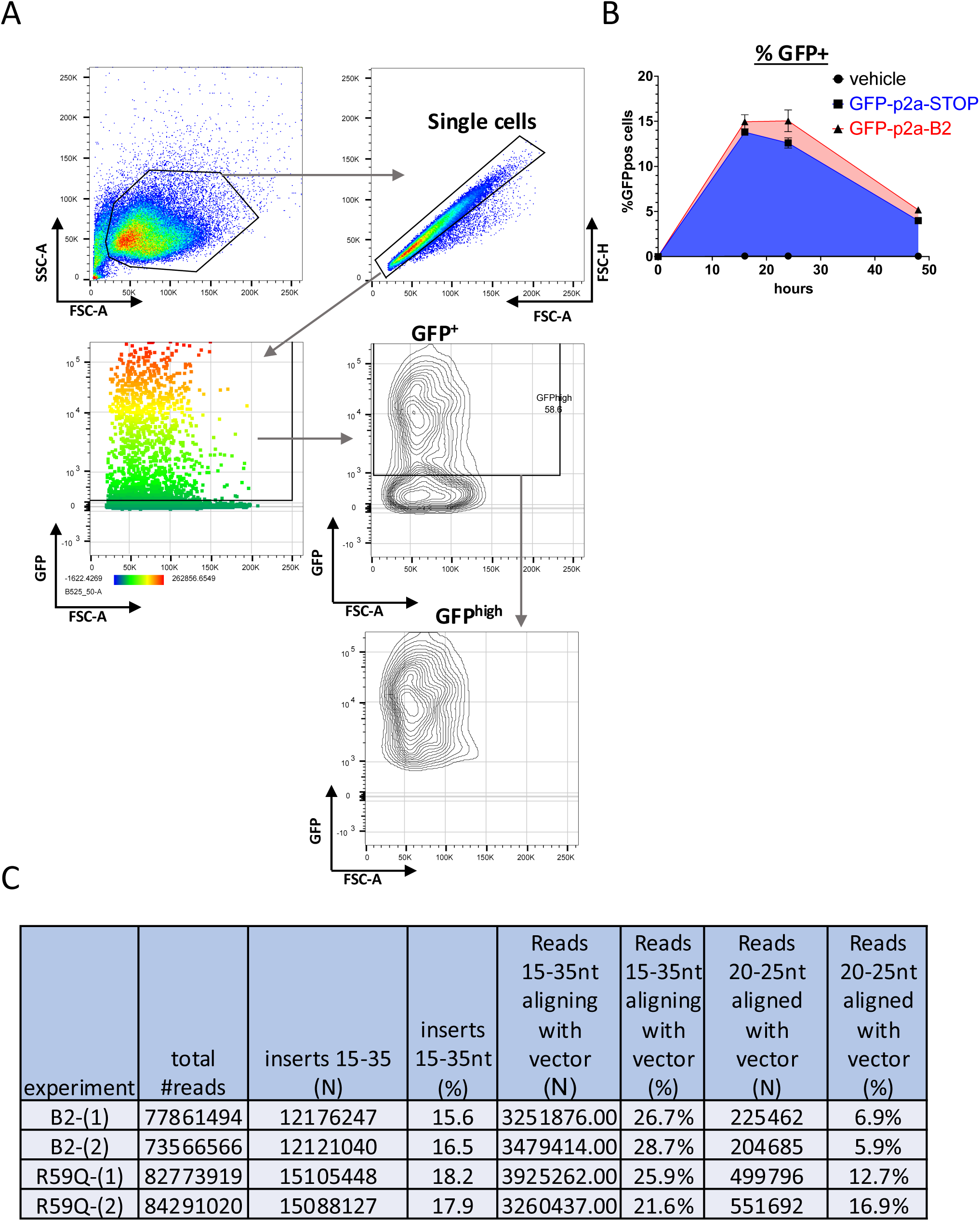
**(A)** Gating strategy used for the flow cytometry-based assay. Cells were first selected based on side scatter area (SSC-A) versus forward scatter area (FSC-A), followed by gating on singlets using FSC-Height (FSC-H) versus FSC-A to exclude doublets. Single cells were then analysed for GFP expression. Cells with a MFI greater than 10³ were classified as GFP^high^. **(B)** Percentage of GFP-positive cells in E14 mESCs transfected with CHIKV-GFP-STOP (blue) or CHIKV-B2^wt^ (red) sa-RNA at 16, 24, and 48 hours post-transfection. **(C)** Summary of small RNA deep sequencing reads. The table displays the number of raw and mapped reads obtained for each sample. Mapping was performed against each corresponding CHIKV-based sa-RNA genome.

**Extended Data Figure 2.**
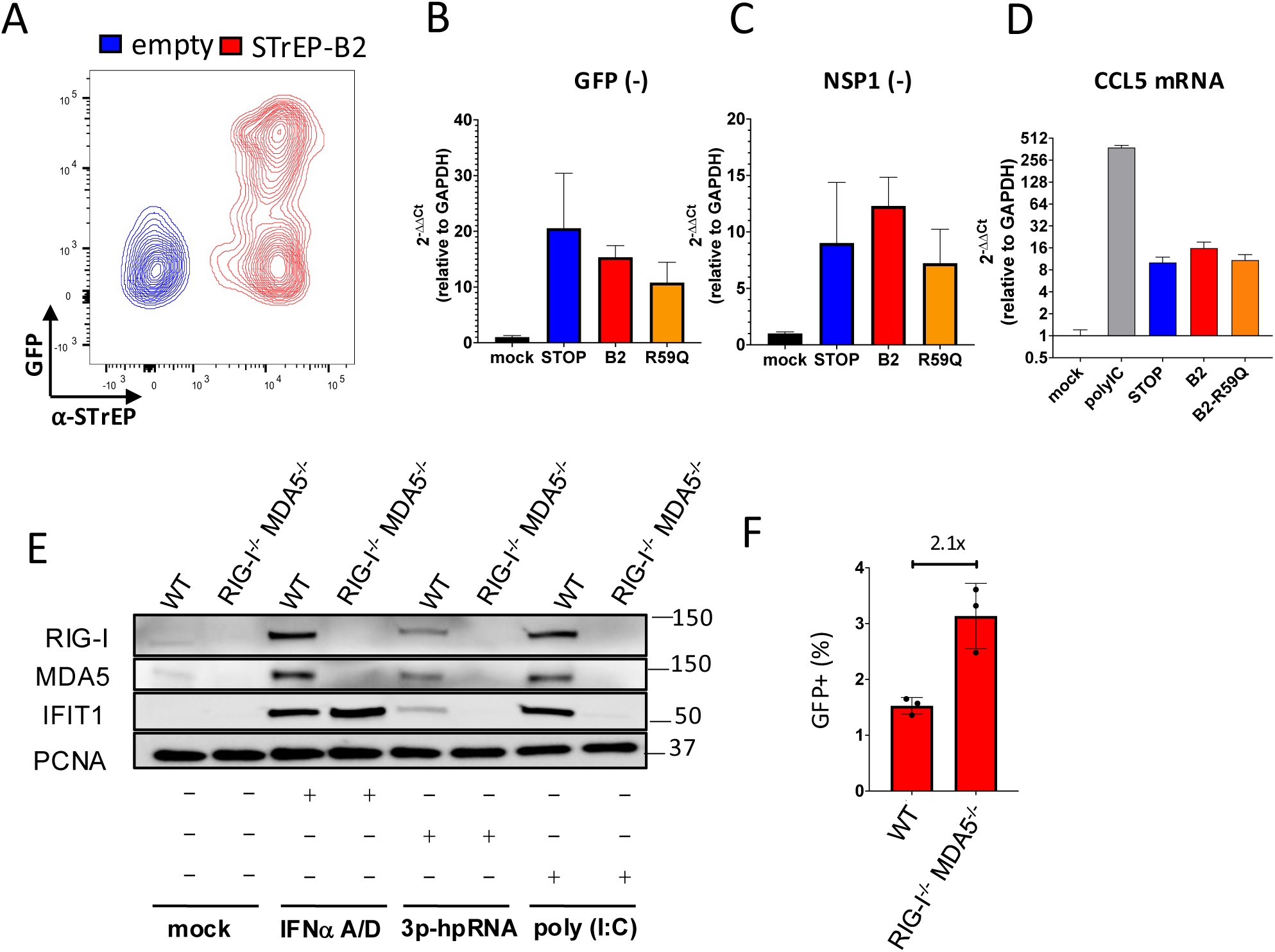
**(A)** HeLa cells were co-transfected with CHIKV-GFP-STOP sa-RNA and either a control vector or a pCAGG construct encoding 1xSTrEP-tagged B2. 24 hours post-transfection, intracellular B2 expression was detected by staining with an anti-STrEP antibody. Cells were gated based on STrEP signal and GFP fluorescence. **(B, C)** HeLa cells were transfected with STOP (blue), B2^wt^ (red) or B2^R59Q^ (yellow) expressing sa-RNA and analysed 24 hours later by RT-qPCR for CHIKV sa-RNA anti-genome levels using primers targeting GFP (B) or CHIKV-nsP1 region (C). **(D)** Cells were treated as in (B,C) and CCL5 mRNA levels was analysed by RT-qPCR **(E)** Western blot analysis of HEK293 WT or RIG-I^-/-^ MDA5^-/-^ cells treated with recombinant universal IFN-I (IFNα-A/D) or transfected with 5′-triphosphate hairpin RNA (3p-hpRNA) or poly (I:C). **(F)** Quantification of GFP^+^ HEK293 WT or RIG-I^-/-^ MDA5^-/-^ cells transfected with CHIKV-GFP-B2^wt^ and analysed by flow cytometry.

**Extended Data Figure 3.**
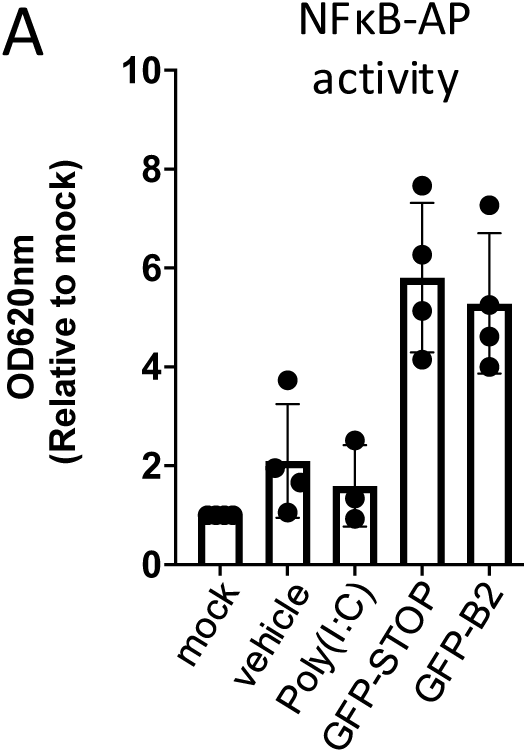
**(A)** THP1-2R-wt cells were transfected with CHIKV-GFP-STOP or -B2^wt^. NF-κB activation was quantified by measuring secreted alkaline phosphatase (SEAP) activity in the supernatant. Data are from 4 independent experiments where each point represents the average of technical replicates.

**Extended Data Figure 4.**
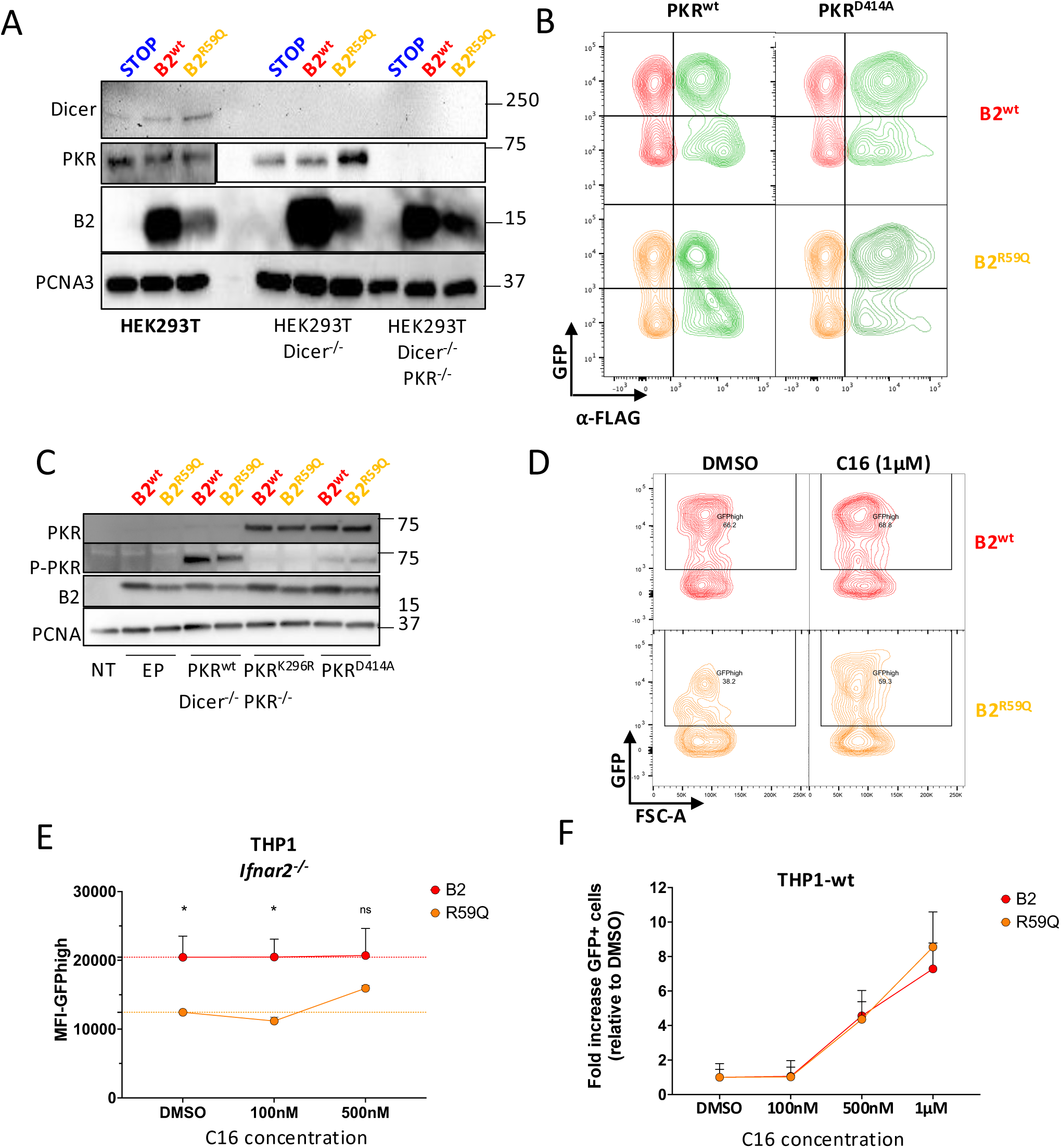
**(A)** Western blot analysis of HEK293T-WT, -Dicer^-/-^, Dicer^-/-^PKR^-/-^ cells 24 hours after transfection with corresponding CHIKV-GFP sa-RNAs. **(B)** Representative contour plots of HEK293T-Dicer^-/-^PKR^-/-^ transfected first with empty plasmid (EP), FLAG-PKR^wt^ or FLAG-PKR^D414A^ and 24 hours later with the indicated CHIKV-GFP sa-RNAs followed by flow cytometry analysis the next day. Cells were stained for FLAG to identify PKR-expressing cells. In red (B2^wt^) and yellow (B2^R59Q^), contour plots represent cells transfected with empty plasmid while PKR expressing cells are labelled green. **(C)** Western blot analysis of experiment described in (B). **(D)** Representative contour plots of THP1 cells 24 hours after transfection with corresponding CHIKV-GFP saRNAs and treated with PKR inhibitor C16 (1µM) or DMSO. **(E)** THP1-*Ifnar2^-/-^*cells were transfected with CHIKV-GFP-B2^wt^ (red) or -B2^R59Q^ (orange) in the presence of increasing concentration of C16. **(F)** Fold increase of GFP^+^ THP1-wt cells transfected with CHIKV-GFP-B2^wt^ (red) or -B2^R59Q^ (orange) in the presence of increasing concentrations of PKR inhibitor C16. Fold increase was calculated relative to the average percentage GFP positive cells calculated for the DMSO control from 4 independent experiments.

**Extended Data Figure 5.**
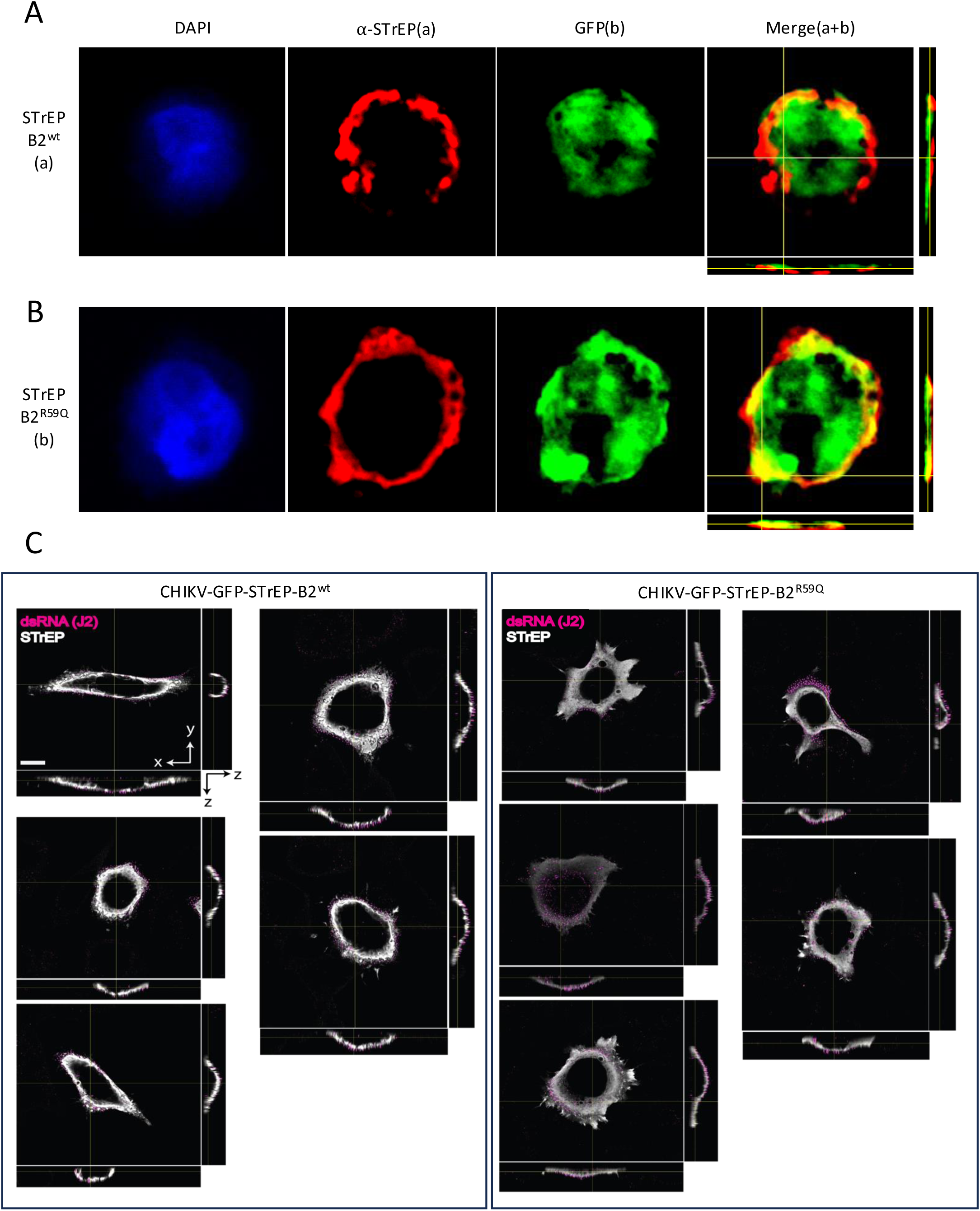
**(A, B)** Confocal microscopy image of THP1-*Ifnar2^-/-^* cells transfected with CHIKV-GFP sa-RNA encoding STrEP-B2^wt^ (A) or STrEP-B2^R59Q^ stained for nucleus (blue) and STrEP-tag (red). **(C)** Hela PKR^-/-^ cells were transfected with CHIKV-GFP sa-RNA encoding STrEP-B2^wt^ (left) or STrEP-B2^R59Q^ (right). Representative images show NoV B2 (STrEP, white) and dsRNA (J2, magenta). Scale bar is 10 um.

**Extended Data Figure 6.**
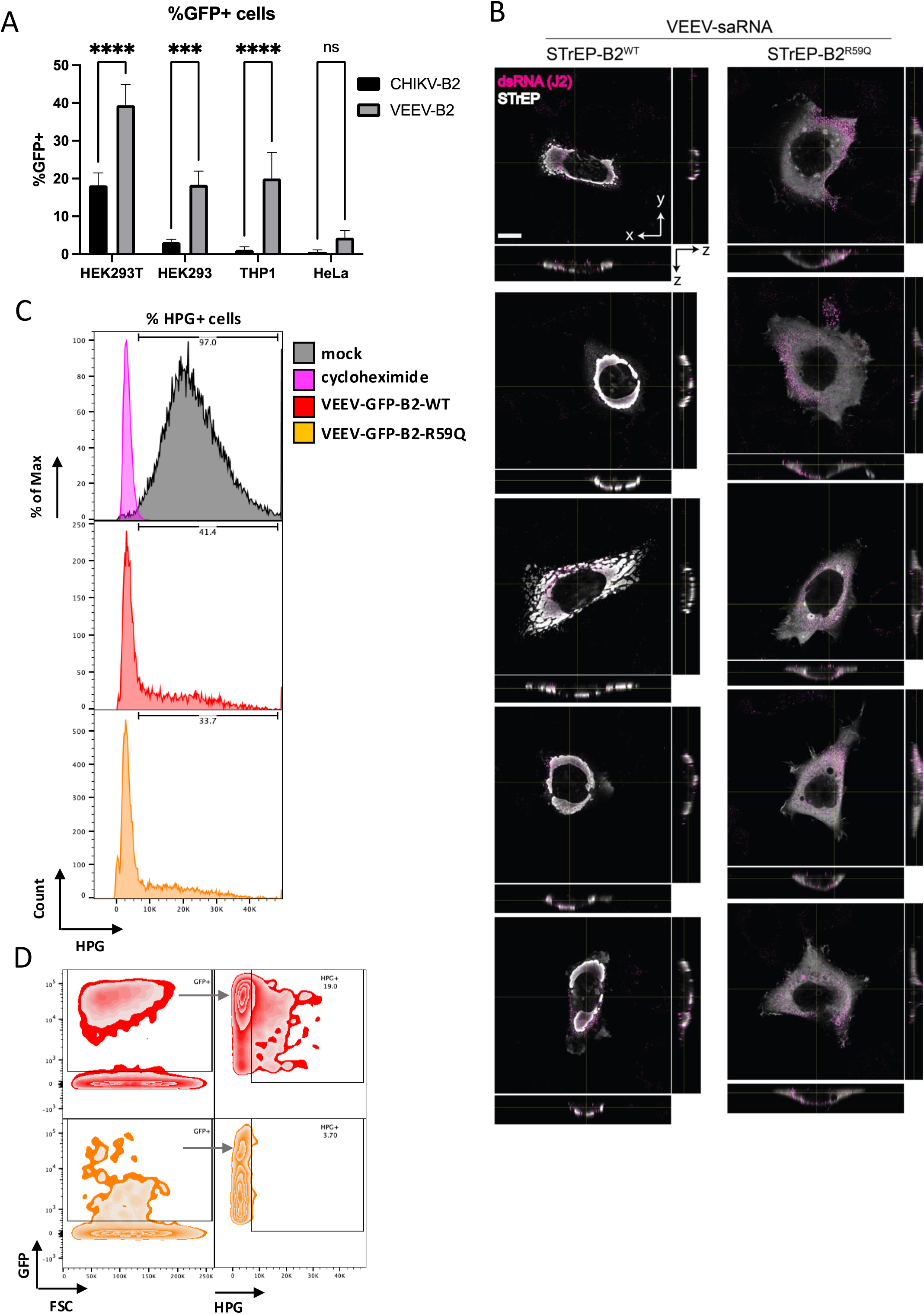
**(A)** Percentage of GFP^+^ cells upon transfection of HEK293T, HEK293, THP1, and HeLa cells using CHIKV-GFP-B2 and VEEV-GFP-B2 sa-RNAs. Data from three independent experiments where each point represents the average of technical replicates is shown. ***P < 0.001, ****P < 0.0001 [two-way ANOVA]. **(B)** Hela cells were transfected with VEEV-GFP sa-RNA encoding STrEP-B2^wt^ (left) or STrEP-B2^R59Q^ (right). Representative images show NoV B2 (STrEP, white) and dsRNA (J2, magenta). Scale bar is 10 um. **(C)** Representative histogram plots of HeLa cells 24 hours after transfection with corresponding VEEV-GFP sa-RNA, followed by HPG incorporation assay in order to measure translation initiation. Samples were compared to mock or cycloheximide-treated cells. **(D)** Gating strategy used for the assessment of translation initiation using HPG click chemistry. Single cells were analysed for GFP expression and HPG incorporation was measured using the Alexa Fluor 594 channel.

## Materials and Methods

### Cloning

CHIKV^ERR^ sa-RNA plasmid was based on replicon vector of CHIKV LR2006OPY1 strain ^89^ a mutation _674_ALT_676_ to _674_ERR_676_ eliminating cytotoxic properties of nsP2 protein ^41^was introduced using site-directed mutagenesis. VEEV sa-RNA plasmid was generated using SP6-VEE-IRES-Puro (Addgene #58971) by introduction of two point mutations in the non-structural region and replacement of IRES-Puro region a multiple cloning site downstream of the subgenomic promoter. Different constructs were generated using standard molecular cloning techniques. Transgene inserts were amplified via PCR/overlap PCR using Q5® High-Fidelity DNA Polymerase (M0491, New England Biolabs), and cloned into CHIKV^ERR^ or VEEV vector using ApaI/PacI restriction sites (New England Biolabs). The GFP sequence present in the engineered constructs is from mGreenLantern (Addgene #161912) ^90^. Codon optimized prefusion SARS-Cov2 Spike-encoding sequence was obtained as synthetic DNA [Genewiz]. Hemaglutinin (HA)-encoding sequence from Influenza PR8 strain was amplified by PCR from a plasmid encoding the wt HA sequence kindly provided by Professor Wendy Barclay (Imperial College London, London, UK)^91^. Backbone plasmids were dephosphorylated after digestion [Antarctic Phosphatase, M0289, New England Biolabs]. Ligation reactions were performed using T4 DNA Ligase [M0202, New England Biolabs] for 16h at 16°C, and bacterial transformation was carried out using NEB 5 alpha competent *E.Coli* cells, a derivative of DH5α [C2987U, New England Biolabs] by heat-shock. Clones were screened by Sanger sequencing and the whole plasmid sequences were verified by Nanopore sequencing technology (Source BioScience). Sequences encoding PKR, B2, and CHIKV-nsPs were amplified by classical PCR/overlap PCR with the corresponding tags added within the forward primer, cloned into pCAGG vector and verified by Sanger sequencing and whole plasmid sequencing. Sequences of all plasmid and primers used for cloning are available upon request.

### sa-RNA in vitro transcription

sa-RNAs were synthesized using mMACHINE™ SP6 Transcription Kit (AM1340, ThermoFisher Scientific) from linearized plasmid DNA template. CHIKV^ERR^ and VEEV plasmids were linearised using NotI (R0189S, New England Biolabs), and MluI (R3198S, New England Biolabs), respectively. Linearised plasmids were run on agarose electrophoresis and gel purified (Monarch DNA Gel extraction kit, New England Biolabs). The *in vitro* transcription reaction was carried out at 37°C for 2 hours with 1µg of template DNA followed by DNase I treatment to remove residual DNA. RNA was purified using silica column-based purification (74106, RNeasy Kit, Qiagen) and quantified using a Nanodrop spectrophotometer (DS-11 DeNovix). RNA integrity was assessed by Bioanalyzer (Agilent 2100).

### Cell culture and cell lines

HeLa (ATCC CCL-2), HeLa-PKR⁻/⁻ (kind gift from Prof. Adam Gaballe), HEK293 (ATCC CRL-1573), and HEK293T (ATCC CRL-3216) cells were maintained in Dulbecco’s Modified Eagle’s Medium (DMEM; 31966, Gibco) supplemented with 10% heat-inactivated fetal bovine serum (FBS, FB-1001, Biosera) and 100 U ml^-1^ penicillin/streptomycin (15140, Gibco) at 37°C in 5% CO₂. THP1-Dual™ (thpd-nfis, InvivoGen) and THP1-Dual™ KO-IFNAR2 cells (InvivoGen) were cultured in Roswell Park Memorial Institute 1640 Medium (RPMI, 21875, Gibco) containing 10% FBS, and100 U ml^-1^ penicillin/streptomycin. E14 mESCs (CRL-1821) were cultured in DMEM supplemented with 10% embryonic stem cell qualified fetal calf serum (FCS, Gibco), 2mM non-essential amino acids (25-025-CIR, Gibco), 2mM sodium pyruvate (S8636, Sigma-Aldrich), 2mM L-glutamine (25030, Gibco), 100 U ml^-1^ penicillin/streptomycin, 0.2% β-mercaptoethanol (31350, Gibco), and 1000 U ml^-1^ ESGRO recombinant mouse leukaemia inhibitory factor (ESG1107, Sigma-Aldrich), at 37°C with 5% CO₂ in gelatine-coated plates. Cells were passaged using trypsin upon reaching 60% confluency. HEK293T Dicer^-/-^ and HEK293T Dicer^-/-^PKR^-/-^ cells were kindly provided by Prof. Bryan R. Cullen (Duke University Medical Center, Durham, USA)^48,49^. HeLa PKR^-/-^ cells were kindly provided by Prof. Adam P. Geballe (Fred Hutchinson Cancer Center, Seattle, USA) ^92^. HEK293 RIG-I^-/-^ MDA5^-/-^ (double KO, DKO) were a kind gift from Dr Annemarthe Van der Veen (Leiden University Medical Center, Netherlands)^93^.

### IFN-I and NF-κB reporter assay with THP1-Dual™ cells

THP1 cells were transfected with saRNAs and, 24 hours later, supernatants (20 μL) were transferred to a white (opaque) 96-well plate (3922, Corning). Secreted Lucia luciferase activity was then measured using the *Renilla* Luciferase Assay System (E2820, Promega) following the manufacturer’s instructions. Luminescence was recorded using a FLUOstar Omega plate reader (BMG LABTECH). Secreted embryonic alkaline phosphatase (SEAP) activity in culture supernatants was quantified using the QUANTI-Blue™ Solution (rep-qbs, InvivoGen). Supernatants were incubated with QUANTI-Blue™ at 37°C for 1h, and absorbance was measured at 620–655 nm using a spectrophotometer.

### sa-RNA transfection and drug treatment

Cells were collected, counted, and resuspended in serum-free Opti-MEM™ before transfection. Transfection complexes were prepared by incubating 200 ng of sa-RNA per 100,000 cells or 1 µg per 500,000 cells with Lipofectamine™ MessengerMAX™ (LMRNA001, Invitrogen) in Opti-MEM™ (31985070, Gibco) for 10–15 min at room temperature. Complexes were added directly to the cell suspension, followed by 30 min incubation at 37°C, 5% CO₂. After transfection, cells were transferred to the appropriate well format with complete medium or, where indicated, complete medium containing PKR inhibitor C16 (527450, Sigma-Aldrich), or 500nM of JAK inhibitor Ruxolitinib (tlrl-rux, Invivogen), or 500 UI ml^-1^ of universal type I IFN (human IFN alpah A/D) (11200, PBL assay Science).

### Fluorescence-Activated Cell Sorting (FACS) and flow cytometry analysis

Cells were harvested by trypsinisation, washed with PBS containing 2% FCS and resuspended in PBS-4%-PFA (40-7401-05, Severn Biotech Ltd.), washed and analysed using an BD LSR II Flow Cytometer acquiring a sample size of 10,000 cells. For samples that required intracellular staining, cells were permeabilized (BD Perm/Wash™, 554723, BD Biosciences), blocked and stained with either a) anti-Flag, b) anti-STrEP, c) anti phospho-eIF2⍺ antibodies for 2 hours at room temperature. Samples were then incubated with secondary antibody anti-mouse AF647 (A-21239, Invitrogen) for 2 hours at room temperature in the dark. Cells were washed 3 times with PBS and resuspended in 200 µl of FACS buffer (PBS, 2% FCS, 2 mM EDTA). Data analysis was performed with FlowJo (TreeStar).

### RNA extraction and quantification

Total RNA was extracted using the RNeasy Mini Kit (Qiagen) / TRIzol reagent (Invitrogen) following the manufacturer’s protocol. Column purification was followed by DNase-treatment according to the manufacturer’s instructions (Qiagen, 79254). RNA concentration and purity were assessed using a Nanodrop spectrophotometer (DS-11 DeNovix) and integrity was confirmed using a Bioanalyzer (Agilent 2100) and agarose gel electrophoresis. Reverse transcription was performed using the SuperScript™ III First-Strand Synthesis System for RT-PCR (18080-051, Invitrogen) with random hexamers or oligo-dT primers. qPCR reactions were set up using SYBR Green Master Mix (4309155, Applied Biosystems) and amplification was performed on a LightCycler 480 System (Roche). Relative expression levels were determined using the ΔΔCt method, normalizing to GAPDH. The sequences of primers used are provided in Supplementary Table S2.

### Small RNA deep sequencing after cell sorting

50’000 GFP positive E14 cells were sorted using a BD FACSAria (BD Biosciences) and collected in FCS-containing medium. Sorted cells were centrifuged and lysed in TRIzol (10296010, Ambion). RNA was extracted using the phenol/chloroform protocol. Experiments were done on biological duplicates, Small RNA deep sequencing was carried out by Fasteris (Genesupport SA), Geneva, Switzerland, Briefly, RNA processing and library preparation were done in parallel. Library preparation for small RNA sequencing was performed using the QiaSeq Small RNA Library Prep Kit, following the manufacturer’s protocol. Sequencing was performed on an Illumina NextSeq sequencer at 1×75 bp read length The adapter sequences were trimmed from the raw reads using the UMI-tools. Inserts of s 15 to 35 bases were selected and mapped with BWA to mouse miRNAs (miRbase-r22_1). Mapped reads counts were used to quantify miRNA expression. Differential expression analysis was conducted using DESeq2.

### Confocal microscopy and image analysis

Cells were seeded onto an ibidi 8-well (80826, ibidi GmbH) chambered coverslip and fixed with 4% paraformaldehyde for 30 minutes at room temperature. Cells were washed with PBS and then permeabilized using 0.5% Triton X-100 in PBS for 30 minutes. The coverslip was then blocked for 2 hours at room temperature with Maxblock (15252, Active Motif) and subsequently incubated with primary antibodies in 0.1% Triton X-100 in PBS overnight at 4°C. Antibody references are provided in Supplementary Table S1. Primary antibodies include mouse anti-strep (1/500), rabbit anti-strep (1/500), rabbit anti-CHIKV-nsP1 (1/200), rabbit anti-CHIKV-nsP2 (1/200), mouse anti-dsRNA J2 (1/200) and rabbit anti-dsRNA J2 (1/200). Before incubation with secondary antibodies (goat anti-rabbit IgG Alexa Fluor™ 405 (1/500), goat anti-mouse IgG Alexa Fluor™ 405 (1/500), goat anti-mouse IgG1 Alexa Fluor™ 568 (1/500), goat Anti-mouse IgG, Alexa Fluor™ 647 (1/500) or goat Anti-rabbit IgG, Alexa Fluor™ 647 (1/500) for 2 hours at room temperature, samples were washed 3 times for 5 minutes with 0.1% Triton X-100. The nuclei were stained with either DAPI (1/1000, 564907, BD Pharmingen) or NucSpot 568/580 (1/5000, 41036, Biotium). In case the nuclei were stained with DAPI, the dye was included in the secondary antibody mix. When an Alexa Fluor 405-conjugated secondary antibody was used, the staining protocol was adapted to include the NucSpot 568/580 which was applied after secondary antibody staining followed by incubation for 5 minutes at room temperature. Next, samples were washed 3 times for 5 minutes with PBS and then preserved in excess of PBS until imaging. Images were acquired using a Zeiss LSM 880 Airyscan confocal microscope. Before image analysis, acquired images are processed using the ‘Airyscan processing’ package in the Zen black (Zeiss) software. Further analysis was done using Arivis (Zeiss) and Image J (Fiji).

Image analysis of dsRNA aggregates was done in Arivis (Zeiss). To that end all images were converted using the plug in the Arivis software (Arivis SIS converter). The analysis consisted of 4 steps: identifying the cells, identifying the dsRNA aggregates, allocating dsRNA to the cells and measuring distance to its centre. Transfected cells were identified using the GFP signal. This signal was first denoised (method; median, diameter; 0.5 μm), and subsequently filtered using intensity (method; simple, object type; bright, threshold depending on the experiment) and size (10-5000 μm^2^). The dsRNA aggregates were identified using the Blob finder (diameter; 0.49 μm, Probability threshold; 14%, Split sensitivity 90%) and then filtered on size (voxel size of 35). Allocating the dsRNA to the cells was done using the compartmentalization tool. All dsRNA aggregates within the GFP, on the border of the GFP or withing 0.49 μm of the border were included. Finally, distances of the dsRNA aggregates to the centre of the GFP were calculated (Distance tool; parent, centre of geometry).

### Nascent protein synthesis assay

HeLa cells were transfected with VEEV-GFP-sa-RNA and, after 20 hours, were washed with PBS and incubated 1 hour in methionine-free media (21013, Gibco) supplemented with L-cystein (C6852, Sigma) to starve the cells prior to priming. Cells were then incubated with L-homopropargylglycine (HPG) to label newly synthesized proteins using the Click-iT™ HPG Alexa Fluor™ 594 Protein Synthesis Assay Kit (C10429, Invitrogen) according to the manufacturer’s instructions. Cells were then trypsinised, fixed with PBS containing 4% paraformaldehyde and HPG incorporation was analysed by flow cytometry.

### Protein co-immunoprecipitation

4 × 10⁶ HEK293T cells were plated in 100 mm dishes and transfected the following day using Lipofectamine 2000 (11668-019, ThermoFisher) with plasmids encoding N-terminal 1xSTrEP -B2, -mCherry, or -B2-R59Q, alongside plasmids encoding CHIKV replication complex components (i.e. nsP1, nsP2^ERR^, nsP3, nsP4) together or independently. Transfections were performed according to the manufacturer’s recommendations and transfected cells were subsequently incubated for 24 hours at 37°C, 5% CO₂. Cells were then lysed in 1 ml of MOPS lysis buffer (20 mM MOPS-KOH pH 7.4, 150 mM KCl, 0.5% NP-40, 2 mM β-mercaptoethanol) supplemented with Complete Mini EDTA-free protease inhibitor cocktail (11836170001, Roche) for 30 minutes on ice, followed by centrifugation (12,000 × g, 15 minutes, 4°C). 50 µl of lysate was collected for total protein samples, remaining lysate was incubated with Strep-Tactin beads (2-1613-002, IBA Lifesciences) overnight at 4°C in a spinning wheel. Beads were washed 4 times with lysis buffer and eluted in 2×50µl of 1× Buffer E (2-1000-025, IBA Lifesciences). Total and eluted protein samples were resolved on a Tris-Glycin SDS-polyacrylamide gel and analysed by Western blot.

### Western Blot

Cells were lysed in RIPA buffer (89900, Thermo Scientific) supplemented with protease (11836170001, Roche) and phosphatase inhibitors (78446, Thermo Scientific). Proteins were resolved on 7.5% or 4-15% or 4-20% Mini-PROTEAN TGX gels (4561023, 4561095, 4561086, BIORAD) and transferred by electroblotting to PVDF membranes (Millipore). Membranes were blocked in either TBS with 0.1% Tween-20 and 5% non-fat milk or 5% milk/TBST (except for detection of phosphorylated residues, at this case 5% BSA was used), incubated with primary antibodies (Supplementary Table S2) overnight at 4°C, followed by incubation with HRP-conjugated secondary antibodies (1:10000, ref). Detection was performed using ECL Western blotting reagents (RPN2232, Cytiva) or SuperSignal West Femto maximum sensitivity substrate (34095, ThermoFisher Scientific). Anti-B2 antibody was kindly provided by Prof. Shou-Wei Ding (University of California, Riverside, USA).

---

**Supplementary Table S1.** (antibodies)

| antibody | company | Cat# |
| --- | --- | --- |
| mouse-anti-Flag | Sigma-Aldrich | F1804 |
| mouse-anti-STrEP | GeneTex | GTX628899 |
| rabbit-anti-STrEP | Novus | NBP2-41075 |
| RIG-I | Cell Signaling | 3747S |
| MDA5 | Cell Signaling | 5321S |
| IFIT1 | Cell Signaling | 14769S |
| PKR | Abcam | AB184257 |
| Phospho-PKR | Abcam | AB32036 |
| eiF2α | Thermo Scientific | AHO0802 |
| Phospho-eiF2α | Abcam | AB32157 |
| vinculin | Abcam | AB129002 |
| CHIKV-NSP1 | Novus | NBP313441 |
| CHIKV-NSP2 | Life Technologies | MA547042 |
| HA-PR8 | Thermo Fischer | PA5-34929 |
| Spike-SARS | Thermo Fischer | PA1-41165 |
| mouse anti-dsRNA J2 | exalpha/nordic | 10010200 |
| rabbit anti-dsRNA J2 | Absolute antibodies | Ab01299-23.0 |
| PCNA | Cell Signaling | 13110S |
| anti-rabbit-HRP | Cell Signaling | 7074S |
| anti-mouse-HRP | Cell Signaling | 7076S |
| Goat anti-Rabbit IgG<br>Alexa Fluor™ 405 | Invitrogen | A48258 |
| Goat anti-mouse IgG<br>Alexa Fluor™ 405 | Invitrogen | A31553 |
| Goat anti-mouse IgG1<br>Alexa Fluor™ 568 | Invitrogen | A21124 |
| Goat Anti-mouse IgG,<br>Alexa Fluor™ 647 | Invitrogen | A32728 |
| Goat Anti-rabbit IgG,<br>Alexa Fluor™ 647 | Invitrogen | A32733 |

**Supplementary Table S2.**
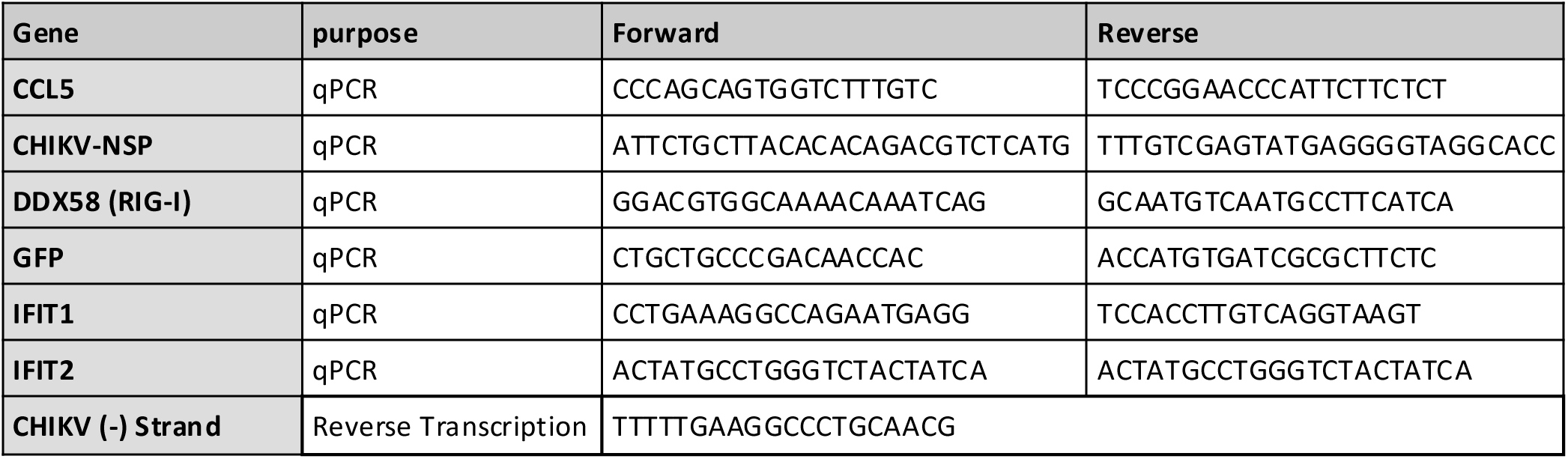
(primers)

